# Accurate estimation of SNP-heritability from biobank-scale data irrespective of genetic architecture

**DOI:** 10.1101/526855

**Authors:** Kangcheng Hou, Kathryn S. Burch, Arunabha Majumdar, Huwenbo Shi, Nicholas Mancuso, Yue Wu, Sriram Sankararaman, Bogdan Pasaniuc

## Abstract

The proportion of phenotypic variance attributable to the additive effects of a given set of genotyped SNPs (i.e. SNP-heritability) is a fundamental quantity in the study of complex traits. Recent works have shown that existing methods to estimate genome-wide SNP-heritability often yield biases when their assumptions are violated. While various approaches have been proposed to account for frequency- and LD-dependent genetic architectures, it remains unclear which estimates of SNP-heritability reported in the literature are reliable. Here we show that genome-wide SNP-heritability can be accurately estimated from biobank-scale data irrespective of the underlying genetic architecture of the trait, without specifying a heritability model or partitioning SNPs by minor allele frequency and/or LD. We use theoretical justifications coupled with extensive simulations starting from real genotypes from the UK Biobank (*N* = 337K) to show that, unlike existing methods, our closed-form estimator for SNP-heritability is highly accurate across a wide range of architectures. We provide estimates of SNP-heritability for 22 complex traits and diseases in the UK Biobank and show that, consistent with our results in simulations, existing biobank-scale methods yield estimates up to 30% different from our theoretically-justified approach.

## Introduction

SNP-heritability, the proportion of phenotypic variance attributable to the additive effects of a given set of SNPs, is a fundamental quantity in the study of complex traits^1^; it provides an upper bound on risk prediction from a linear model relating genotypes to phenotype^2^ and, when defined as a function of all SNPs on a genotyping array, yields insights into the “missing heritability” of complex traits^3–5^. Traditionally, SNP-heritability is estimated by fitting variance components models with REML^3,6–9^. With some notable exceptions^8^, REML-based methods are typically not scalable to biobanks that assay hundreds of thousands of individuals (e.g., UK Biobank contains genotype measurements for more than half a million individuals^10^). SNP-heritability can also be estimated from summary-level GWAS data by assessing the deviation in marginal association statistics as a function of the LD score of each SNP^11–14^, thus making SNP-heritability estimation scalable to hundreds of thousands or even millions of individuals. More recently, a randomized extension of Haseman-Elston (HE) regression^15^ was shown to estimate a single genetic variance component from individual-level data as accurately as REML methods but in a fraction of the run-time^16^.

To facilitate inference, all existing methods for genome-wide SNP-heritability inference make various assumptions on the underlying genetic architecture of the trait, which is typically parametrized by *polygenicity* (the number of variants with effect sizes larger than some small constant δ) and *MAF/LD-dependence* (the coupling of effect sizes with minor allele frequency (MAF), local linkage disequilibrium (LD), or other functional genomic annotations such as regions of open chromatin)^17^. Since the true genetic architecture of any given trait is unknown, existing methods are susceptible to bias and often yield vastly different estimates of SNP-heritability for the same traits, even when applied to the same data^9,14,18^. Although multi-component methods that stratify SNPs by MAF and LD can ameliorate some of the robustness issues of single-component methods^7,18,19^, fitting multiple variance components to biobank-scale data with REML is highly resource-intensive^8^ and it is currently unclear whether stratifying by MAF/LD produces accurate estimates of total SNP-heritability for methods based on summary statistics. Alternate methods explicitly model MAF- and LD-dependent architectures when estimating SNP-heritability^6,9,14^; however, these approaches can produce drastically different estimates when their assumptions are violated^6,9,14,18,19^. In addition, genetic architecture is unlikely to be the same across traits or populations due to, for example, variable degrees of negative selection acting on different traits in different populations^17,20–25^. Methods that jointly infer SNP-heritability and other parameters such as the strength of negative selection or polygenicity have been proposed^14,23,26^ but are computationally intensive and/or sensitive to LD-dependent architectures. Thus, it remains unclear which estimates of genome-wide SNP-heritability computed from biobank-scale data (e.g., UK Biobank^10^) are reliable.

In this work, we investigate whether genome-wide SNP-heritability can be accurately estimated under a generalized random effects (GRE) model that makes minimal assumptions on the genetic architecture of complex traits. Under this model, every causal effect can have an arbitrary SNP-specific variance, and SNP-heritability is defined as the sum of the SNP-specific variances (Methods). To the best of our knowledge, all existing methods make additional assumptions on top of the GRE model (Table 1). For example, the infinitesimal model assumed by single-component GREML^3^ (and several other methods^8,16,27^) imposes an inverse relationship between MAF and effect size by assuming that every standardized effect size explains an equal portion of total SNP-heritability, whereas the single-component LDAK model assumes that each SNP-specific variance is inversely proportional to both MAF and the LD neighborhood of the SNP^6,9^. We derive a closed-form estimator for SNP-heritability as a function of GWAS marginal association statistics and in-sample LD and show that this estimator is consistent (i.e. approaches the true SNP-heritability as sample size increases) and unbiased (i.e. its expectation is equal to the true SNP-heritability) when the number of individuals is larger than the number of SNPs. Most importantly, the accuracy of this estimator does not depend on the underlying genetic architecture of the trait. While the GRE estimator is similar in form to previously proposed “fixed effect estimators,”^28,29^ our approach differs from previous work in two main ways. First, SNP-heritability defined under a fixed effect model is different from the estimand of interest here (Methods). Second, previous work applied the estimator locally to identify regions that contribute disproportionately to the genome-wide signal^28,29^; in this work, we define a different genome-wide estimator (Equation 1) that requires large-scale genotype data. In addition, previous work applied an SVD-based regularization to introduce bias in favor of reduced variance^29^ whereas in this work, the regularization was unnecessary (all LD matrices used are full rank; see Methods).

**Table 1.** Existing methods to estimate SNP-heritability impose additional assumptions on top of the generalized random effects (GRE) model. Under the GRE model, the causal effects at any two SNPs are assumed to be independent (E[*β*_*i*_*β*_*j*_] = 0 for all *i* ≠ *j*) and genome-wide SNP-heritability is defined as 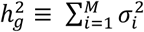, where each 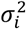 can be an arbitrary nonnegative real number as long as 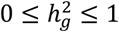 (Methods). All existing methods make assumptions on the distribution of *β*_*i*_ and/or the form of 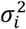 that can be subsumed under the GRE model. To simplify notation, we assume for each model that phenotypes are standardized in the population (i.e. Var[*y*_*n*_] = 1 for every individual *n*).

| Model | Assumptions on $\beta_i$ | Description |
| --- | --- | --- |
| Generalized random effects | $E[\beta_i] = 0, \text{Var}[\beta_i] = \sigma_i^2, \sigma_i^2 \geq 0$ | Each SNP $i$ has a nonnegative SNP-specific variance $\sigma_i^2$. Total SNP-heritability is $h_g^2 \equiv \sum_{i=1}^M \sigma_i^2$. |
| GREML-SC<sup>3,8,16</sup> | $\beta_i \sim N(0, h_g^2/M)$ | Each SNP explains an equal portion of $h_g^2$. In other words, $\sigma_i^2 = h_g^2/M$ for all $i = 1, \dots, M$. |
| GREML-MC<sup>7,8,18,46,47</sup> | $\beta_i \sim N(0, \sum_{c \in \mathcal{C}} [\text{SNP}_i \in c] h_c^2/m_c)$ | $h_g^2$ is partitioned by a set of disjoint SNP partitions $\mathcal{C}$ that span all $M$ SNPs. Partition $c \in \mathcal{C}$ contains $m_c$ SNPs that have per-SNP variances $h_c^2/m_c$. Total SNP-heritability is $h_g^2 = \sum_{c \in \mathcal{C}} h_c^2$. |
| LDAK<sup>6,9</sup> | $\beta_i \sim N(0, \sigma_i^2), \sigma_i^2 \propto w_i [f_i(1 - f_i)]^{1+\alpha}$ | Each SNP-specific variance is proportional to a function of $f_i$ (the MAF of SNP $i$) and to $w_i$ (a SNP-specific weight that is a function of the inverse of the LD score of SNP $i$). $\alpha$ controls the relationship between $\sigma_i^2$ and $f_i$. The most recent recommendation by ref.<sup>9</sup> is to assume $\alpha = -0.25$. |
| LDSC<sup>11</sup> | $E[\beta_i] = 0, \text{Var}[\beta_i] = h_g^2/M$ | Each SNP explains an equal portion of $h_g^2$ (similar to the GREML-SC model when $h_g^2$ is defined with respect to the same set of $M$ SNPs). |
| S-LDSC<sup>12,13,30</sup> | $E[\beta_i] = 0, \text{Var}[\beta_i] = \sum_{a \in A} \tau_a a(i)$ | Each SNP-specific variance is a linear function of a set of annotations $A$ where each $a \in A$ represents a binary or continuous-valued annotation. $a(i)$ is the value of annotation $a$ at SNP $i$. $\tau_a$ is the expected contribution of a one-unit increase in annotation $a$ to each SNP-specific variance. |
| SumHer<sup>14</sup> | $E[\beta_i] = 0, \text{Var}[\beta_i] \propto w_i [f_i(1 - f_i)]^{1+\alpha}$ | An extension of the LDAK model to operate on summary-level data; can also efficiently partition $h_g^2$ by multiple annotations. The most recent recommendations by refs.<sup>9,14</sup> is to set $\alpha = -0.25$. |

Through theoretical derivations and extensive simulations across a wide range of MAF- and LD-dependent architectures starting from real genotypes from the UK Biobank^10^ (337K individuals and 593K SNPs), we find that the GRE estimator provides nearly unbiased estimates of SNP-heritability across all architectures whereas existing methods are sensitive to model misspecification. For example, across 126 distinct architectures, the maximum bias we observe with the GRE estimator is 2% of the simulated SNP-heritability whereas methods such as stratified LD score regression (S-LDSC)^12,13^ and SumHer^14^ yield biases between −64% and 28%. For completeness, we also contrast the GRE estimator with several REML-based methods in simulations at lower sample sizes (due to the computational burden of most REML methods) and find that, consistent with recent reports^18^, all REML-based methods are biased when their model assumptions are violated. Across a similar set of 126 architectures, the bias of the GRE estimator ranges from −5% to 6% of the simulated SNP-heritability whereas single-component REML methods^3,6,8,9^ are biased by anywhere between −44% and 18%. We confirm that multi-component REML methods that stratify SNPs by MAF and LD score (GREML-LDMS-I^18^) are more accurate than single-component REML methods if favorable SNP stratification criteria are used (i.e. if SNPs are stratified by the same MAF bins used to define the causal variant MAF spectrum). The performance of the GRE estimator, which does not stratify SNPs or assume a specific heritability model^6,9,14^, is similar to that of GREML-LDMS-I with favorable stratification criteria, thereby confirming that SNP-heritability can be accurately estimated without knowledge of the underlying genetic architecture.

Finally, we use marginal association statistics and in-sample LD from *N* = 290K unrelated British individuals genotyped at *M* = 460K SNPs (MAF > 1%) to provide estimates of SNP-heritability for 22 complex traits and diseases in the UK Biobank^10^. Consistent with our simulations, across the 18 traits with SNP-heritability estimates greater than 0.05, we find that estimates from S-LDSC (controlling for the baseline-LD model^13^) and SumHer differ from the GRE estimates by a median of −9% and 11%, respectively. For example, for height, estimates from S-LDSC (0.56) and SumHer (0.63) are approximately 7% lower and 5% higher, respectively, than our estimate of 0.60. Similarly, for hypertension, estimates from S-LDSC (0.14) and SumHer (0.18) are ±12.5% different from our estimate of 0.16. Taken together, our results demonstrate that SNP-heritability can be accurately estimated from biobank-scale data without prior knowledge of the genetic architecture the trait, motivating the development of new methods to make inferences from biobank-scale data under fewer modeling assumptions.

## Results

### Overview of the approach

We investigate the utility of an estimator for SNP-heritability derived under a model that makes minimal assumptions on genetic architecture. We assume the standardized phenotype of an individual is a linear function of their genotypes: *y* = **x**^***T***^ ***β*** + *ϵ*, where **x** is a vector of standardized genotypes at *M* SNPs, ***β*** is a vector of standardized effect sizes corresponding to the *M* SNPs, and 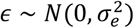 is environmental noise (Methods). We assume the effects can follow any distribution as long as the effect size of every SNP *i* is zero-centered (E[*β*_*i*_] = 0) with a finite SNP-specific variance 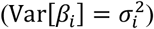 that is allowed to be 0, and that the covariance between the effects of any pair of SNPs is zero (E[*β*_*i*_*β*_*j*_] = 0 for all *i* ≠ *j*). We term this model the “generalized random effects” (GRE) model as, to the best of our knowledge, all existing methods to estimate SNP-heritability impose additional assumptions on top of this model. For example, setting 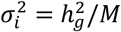 for *i* = 1, …, *M* results in the single-component GREML model^3^, whereas setting 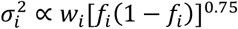 (where *w*_*i*_ is a function of the “LD score” of SNP *i* and *f*_*i*_is the MAF of SNP *i*) results in the most recent LDAK model^9^ (Table 1). Under the GRE model, the SNP-heritability explained by the *M* SNPs is the sum of SNP-specific variances: 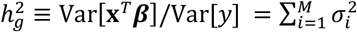 (Methods).

In this work, we are interested in accurately estimating 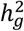 from genotype measurements across *N* individuals at *M* typed SNPs. When *N* > *M*, the estimator 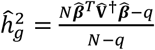, where 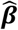 is the vector of standardized SNP effects estimated by ordinary least squares (OLS), 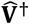 is the pseudoinverse of the in-sample LD matrix, and *q* is the rank of the in-sample LD matrix, is an unbiased estimator of SNP-heritability under the GRE model. That is, 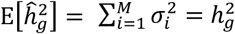 (Methods). The GRE model allows each SNP-specific variance 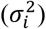 to be an arbitrary finite value satisfying the constraints 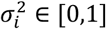 and 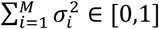. Thus, 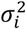 can capture any relationship between effect size and MAF/LD, which in turn implies that 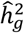 is unbiased under most genetic architectures. Unfortunately, even the largest biobank-scale datasets currently available contain fewer unrelated individuals than typed SNPs (i.e. UK Biobank has genotyped *M* ≈ 593K SNPs in *N* ≈ 337K unrelated British individuals), which limits the utility of the above estimator. We therefore extend our approach by partitioning the genome by chromosome into 22 approximately independent regions:

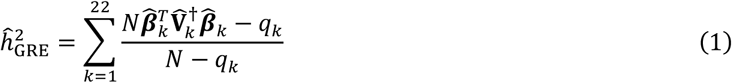

where for each chromosome *k* with *p*_*k*_ typed SNPs, 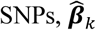 is the *p*_*k*_-vector of standardized SNP effects estimated by ordinary least squares (OLS), 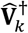 is the pseudoinverse of the in-sample LD matrix, and *q*_*k*_ is the rank of the in-sample LD matrix. Although this genome-wide estimator does not provide theoretical guarantees of unbiasedness, we show through extensive simulations that the magnitude of the bias is extremely small across all architectures when *N* is sufficiently larger than *p*_*k*_.

### Accurate estimation of SNP-heritability irrespective of disease architecture

To investigate the bias and variance of 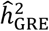, we perform simulations starting from the real genotypes of *N* = 337205 unrelated British individuals in the UK Biobank^10^. First, we use data from chromosome 22 (*M* = 9654 typed SNPs) to simulate 64 distinct MAF- and LD-dependent architectures by varying the SNP-heritability 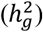, the proportion of causal variants (*p*_casual_), the distribution of causal variant MAF (CV MAF), and the strength of coupling between effect size and MAF/LD; we use “LDAK-LD-dependent” to describe architectures where causal effects are coupled with “LDAK weights” (Methods). To enable comparison of estimates across different values of 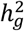, we assess bias as a percentage of the simulated value of 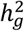 (relative bias) or the error of a single estimate as a percentage of 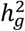 (relative error). Consistent with analytical derivations, the GRE estimator restricted to chromosome 22 provides unbiased estimates across the 64 quantitative trait architectures after correcting for 16 independent tests at each value of 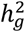 (bias p-value < 0.05/16 is considered significant; see Methods) (Figure 1ac, Supplementary Table S1). The average relative bias across the 64 quantitative trait architectures is 0.00015% of the simulated 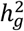, and the largest bias we observe under any single architecture is approximately 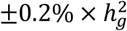 (Supplementary Figure S1a, Supplementary Table S1). In simulations of unascertained case-control studies (Methods), the GRE estimator is approximately unbiased for a range of values of disease prevalence (for 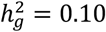, relative bias range is [−0.20%, 0.30%]) and has larger variance for diseases with lower prevalence (Supplementary Figure S2a, Supplementary Table S2). For ascertained case-control studies, estimates are downward-biased but invariant to disease architecture (e.g., when 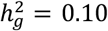, population prevalence = 0.10, and *N*_case_ = *N*_control_, relative bias is approximately −4%) (Supplementary Table S3). We then performed simulations in which 0%, 50%, or 100% of causal SNPs were masked from the observed summary statistics (i.e. untyped). When causal variants are drawn from the MAF range [0.01, 0.05], GRE is downward biased due to lower average LD between the observed typed SNPs and the masked causal SNPs (Supplementary Figure S3). We confirm that the analytical estimator of the standard error (Methods) is well-calibrated across all genetic architectures (Supplementary Figure S4a, Supplementary Table S4). We then investigate the bias induced by partitioning chromosome 22 into non-independent blocks and find that, as expected, our estimator accrues statistically significant upward bias as the average block size decreases (Supplementary Figure S5, Supplementary Table S5). For example, in simulations on chromosome 22 where 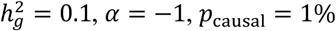, *α* = −1, *p*_casual_ = 1%, and causal variants were drawn uniformly from all SNPs, using a single chromosome-wide LD block produces approximately unbiased estimates (bias = 6.9 × 10^−5^, p-value = 0.55) whereas partitioning the chromosome into 2 disjoint blocks of equal size induces a small but significant upward bias (bias = 4.3 × 10^−4^, p-value = 5.3 × 10^−4^) (Supplementary Figure S5, Supplementary Table S5).

**Figure 1.**
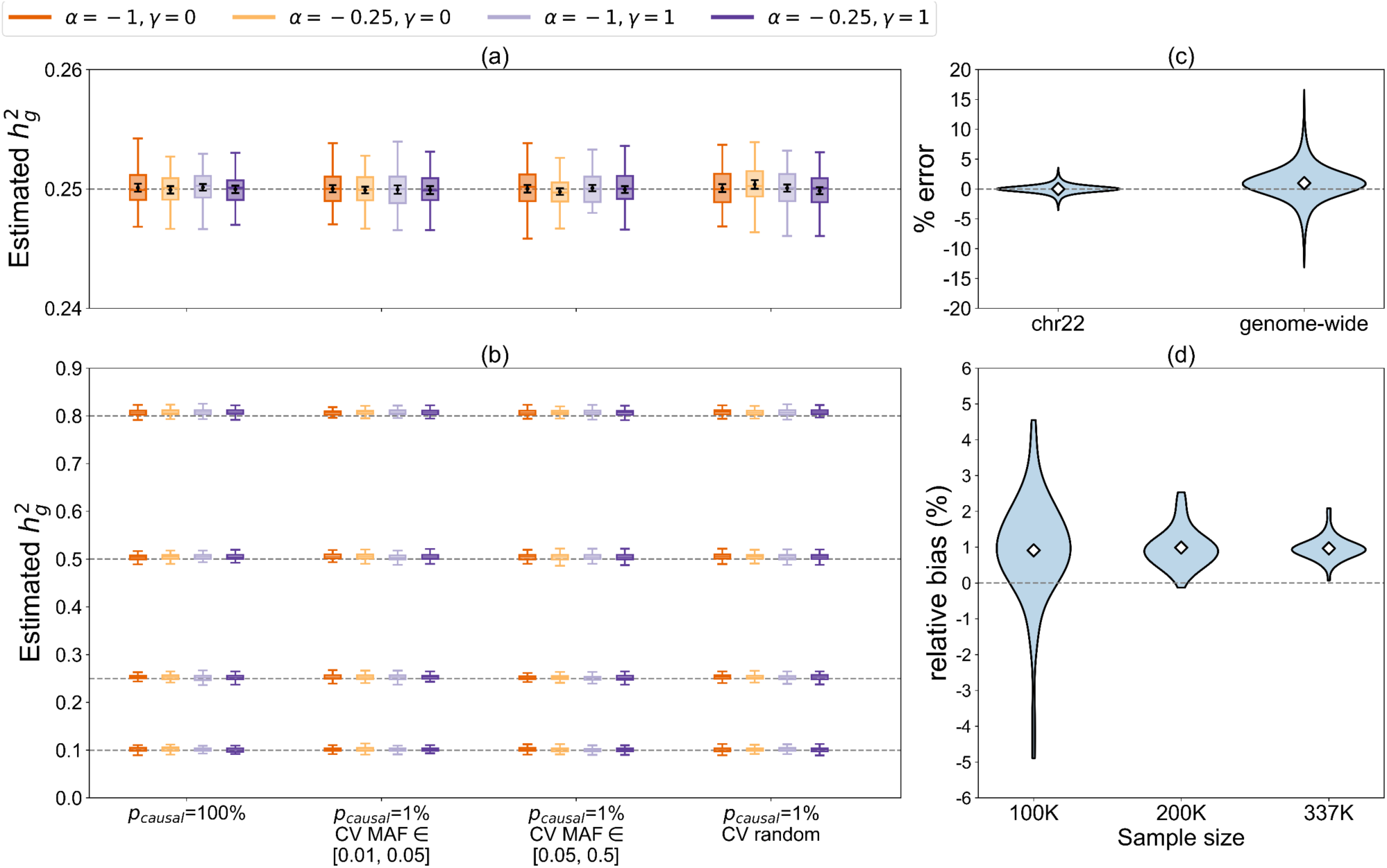
Simulations under 64 distinct MAF- and LD-dependent architectures (*N* = 337205 unrelated British individuals, UK Biobank). For each value of 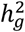, phenotypes were drawn according to one of 16 genetic architectures defined by the polygenicity (*p*_causal_), the MAF range of causal variants (CV MAF), the coupling of MAF with effect size (*α*), and the effect of local LD on effect size (*γ* = 0 indicates no LD weights and *γ* = 1 indicates LDAK weights; see Methods). (a) Distribution of 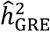 in simulations on chromosome 22 (*M* = 9654 typed SNPs) where 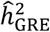 was computed with 1 chromosome-wide LD block. (b) Distribution of 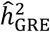 in genome-wide simulations (*M* = 593300 typed SNPs) where 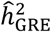 was computed with 22 chromosome-wide LD blocks. In (a) and (b), each boxplot shows the distribution of estimates from 100 simulations. Boxplot whiskers extend to the minimum and maximum estimates located within 1.5×IQR from the first and third quartiles, respectively. Black points and error bars in (a) represent the mean of the distribution and ±2 standard errors of the mean (s.e.m.), which were used to test whether the bias under a single architecture is significant (Methods). (c) Distribution of errors 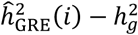, where 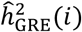 is the estimate from the *i*-th simulation under a given genetic architecture, as a percentage of 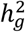. Each violin plot represents the errors of 6400 estimates (64 genetic architectures × 100 simulation replicates). (d) Distribution of relative bias (as a percentage of 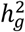) as a function of sample size (*N* = 100K, 200K, or 337K) in genome-wide simulations. Each violin plot represents the distribution of the relative bias of 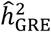 across of 64 genetic architectures. In (c) and (d), the white diamonds mark the mean of each distribution.

Next, we investigate the accuracy of the GRE estimator in genome-wide simulations (*N* = 337K unrelated individuals and *M* = 593K array SNPs) where we use 22 chromosome-wide LD blocks to compute 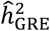. Despite the 22-block approximation, we find that 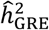 is highly accurate and robust across all 64 MAF- and LDAK-LD-dependent quantitative trait architectures (Figure 1b, 1c). The average bias across the 64 architectures is 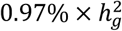, with the relative bias under any single architecture ranging from 0.07% to 2.1% of the simulated 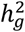 (Supplementary Figure S1b, Supplementary Table S6). The largest error we observe for a single estimate across all 6400 simulations (64 genetic architectures × 100 simulation replicates) is approximately 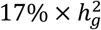 (Figure 1c) and as *N*/*M* increases, the variance of 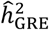 decreases while the relative bias across the 64 architectures appears to be approximately fixed, ranging between 0.91% (*N* = 100K) and 0.99% (*N* = 200K) (Figure 1d). These trends hold for a range of values of *p*_causal_ (Supplementary Figure S6, Supplementary Table S6), for unascertained case-control studies (Supplementary Figure S2b, Supplementary Table S7), and in a smaller set of simulations with *N* = 7685 individuals of South Asian ancestry and *M* = 1642 SNPs (Supplementary Table S8; Methods). Most importantly, the accuracy of the GRE estimator does not correlate with the simulated trait architecture (Figure 1b). We also assess the calibration of our analytical estimator for the standard error in the genome-wide simulations and observe a small downward bias with respect to the empirical standard deviation of 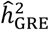 estimates (Supplementary Figure S4b, Supplementary Table S9). For example, across 16 distinct architectures where 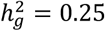, the empirical standard deviation computed from 100 independent estimates of 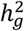 ranges from 0.0049 to 0.0064, whereas our estimate of the standard error is approximately 0.0036 across all architectures (Supplementary Figure S4b, Supplementary Table S9).

We then investigate the effects of unmodeled substructure and/or cryptic relatedness by filtering individuals at different kinship coefficient thresholds (Methods) and find that using stricter relatedness thresholds increases the variance of the estimates (due to smaller sample size) while reducing bias, albeit not significantly (Supplementary Figure S7, Supplementary Table S10). In addition, to assess the impact of population stratification, we simulated an effect of the first genetic principal component (PC) on phenotype and computed OLS association statistics both with and without adjusting for the first genetic PC (Methods). As expected, OLS with no PC adjustment yields inflated estimates while OLS adjusted for the first PC yields approximately unbiased estimates (Supplementary Figure S8, Supplementary Table S11). However, even when a relatively large proportion of phenotypic variance is explained by the first PC (e.g., 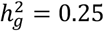 and 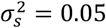), the maximum bias we observe from unadjusted OLS association statistics is 5% of the simulated SNP-heritability (bias p-value = 2.7 × 10^−9^). Together, these results indicate that the GRE estimator is relatively robust to modest amounts of unmodeled substructure and/or population stratification. In all subsequent analyses, we compute 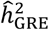 with the 22 chromosome-wide LD block approximation as this provides sufficiently accurate estimates and a fair comparison to other methods.

### Comparison of methods to estimate SNP-heritability

We compare 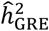 with existing state-of-the-art approaches to estimate SNP-heritability that are easily scalable to the full UK Biobank data (*N* = 337K): LD score regression with no annotations (LDSC), which assumes *α* = −1 and no coupling of effect size with LD^11^; stratified LD score regression (S-LDSC), which partitions 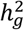 by a set of annotations of interest^12,13^; and SumHer, a recent scalable extension of LDAK which explicitly models MAF- and LD-dependent architectures through a specific form of the SNP-specific variances^14^ (Table 1). To ensure a fair comparison among the methods, LD scores for LDSC, S-LDSC, and SumHer are computed using in-sample LD among the *M* SNPs, and in all simulations we aim to estimate the SNP-heritability explained by the same set of *M* SNPs (see Methods).

We find that 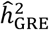 is highly accurate and robust across all simulated architectures while LDSC, S-LDSC, and SumHer are sensitive to deviations from their respective model assumptions. For example, when 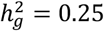 (Figure 2), LDSC is approximately unbiased under the “single-component GREML model” (relative bias = 0.04%, p = 0.86) but is sensitive to the MAF spectrum of causal variants and the degree of coupling between effect size and MAF/LD (e.g., across the 12 architectures where *p*_causal_ = 1%, relative bias ranges from −44% to 50%) (Supplementary Table S12). Similarly, SumHer is accurate under the “LDAK model” (relative bias = 5.3%) but highly sensitive to other plausible genetic architectures (when *p*_causal_ = 1%, relative bias ranges from −19% to 22%) (Figure 2, Supplementary Table S13). Estimates from S-LDSC (MAF), which partitions 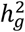 by 10 MAF bins (Supplementary Table S14; Methods), are less biased compared to estimates from LDSC when causal effects are coupled with only MAF, but are significantly downward biased when causal effects are also coupled with LDAK weights (for 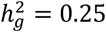, relative bias range is [1.9%, 7.0%] when *γ* = 0 and [−58%, −37%] when *γ* = 1) (Figure 2, Supplementary Table S15). S-LDSC with 10 MAF bins and an additional continuous “level of LD” (LLD) annotation, which we denote S-LDSC (MAF+LLD) (Methods), produces similar results on the same architectures (for 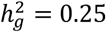, relative bias range is [1.8%, 6.5%] when *γ* = 0 and [−80%, −33%] when *γ* = 1) (Supplementary Table S16). In contrast, the relative bias of 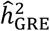 ranges from 0.45% to 1.3% across the same 16 genetic architectures where 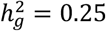 and *p*_causal_ = 1% (Figure 2, Supplementary Table S6). These trends hold for a range of values of 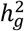 and *p*_causal_: across 112 distinct LDAK-LD- and/or MAF-dependent architectures, the average and range of the relative bias of each method are 0.96% [−0.06%, 2.1%] for 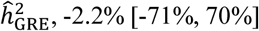, for LDSC, −22% [−62%, 8.7%] for S-LDSC (MAF), −29% [−89%, 9.0%] for S-LDSC (MAF+LLD), and 2.8% [−27%, 28%] for SumHer (Figure 1b, Figure 2, Supplementary Figures S9-S12, Supplementary Tables S6, S12, S13, S15, S16). We also perform simulations under 14 alternative LD-dependent architectures where the variance of each SNP is coupled with its inverse LD score instead of its LDAK weight (i.e. “LD-score-dependent” architectures; see Methods, Supplementary Figure S13) and find that 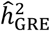 remains nearly unbiased (relative bias ranges from 0.52% to 1.3%) whereas estimates from S-LDSC (MAF), S-LDSC (MAF+LLD), and SumHer are downward-biased on average across the 14 architectures (Supplementary Figure S14, Supplementary Table S17).

**Figure 2.**
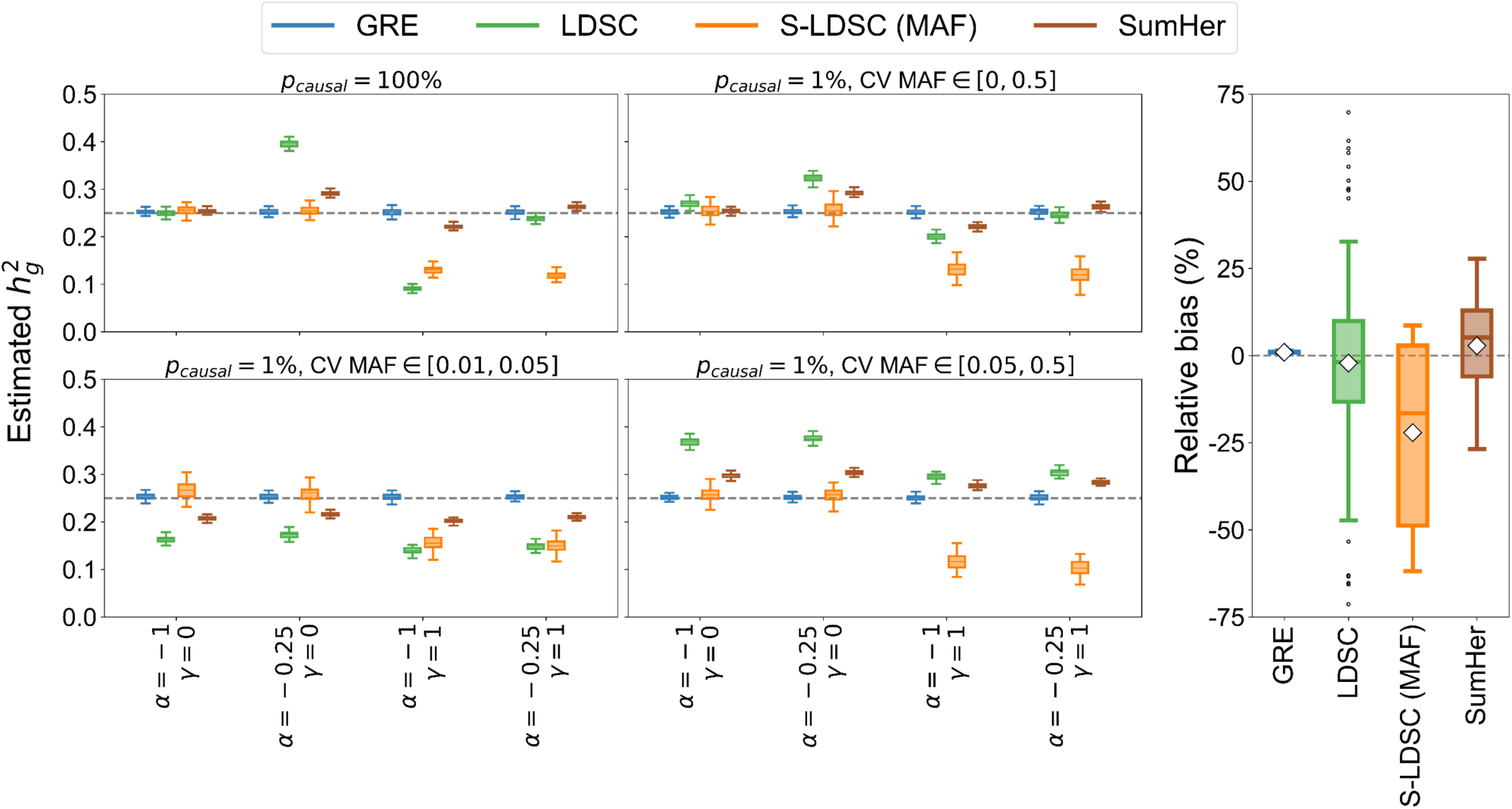
Comparison of 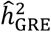 with LDSC, S-LDSC (MAF), and SumHer in genome-wide simulations (*N* = 337205 unrelated individuals, *M* = 593300 array SNPs,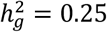). **Left:** Phenotypes were drawn under one of 16 MAF- and/or LDAK-LD-dependent architectures by varying *p*_causal_, *α, γ*, and CV MAF (see Methods). Each boxplot contains estimates of 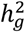 from 100 simulations. Boxplot whiskers extend to the minimum and maximum estimates located within 1.5 × IQR from the first and third quartiles, respectively. **Right:** Relative bias of each method (as a percentage of the true 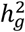) across 112 distinct MAF- and LDAK-LD-dependent architectures (see Methods). Each boxplot contains 112 points; each point represents the average estimated 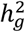 from 100 simulations under a single genetic architecture. The white diamonds mark the average of each distribution.

For completeness, we also compare to four widely used REML-based methods: single-component GREML (GREML), which assumes *α* = −1 and no coupling of effect size with LD^3^; GREML-LDMS-I, a multi-component extension of GREML that partitions SNPs by MAF and LD score^18^; BOLT-REML, a computationally efficient variance components estimation method with assumptions similar to those of GREML^8^; and LDAK, which assumes a specific form of the coupling of effect size with LD and recommends setting *α* = −0.25 ^6,9^ (Table 1). Because it is computationally intractable to apply the REML-based methods to thousands of genome-wide simulations with 337K individuals, we perform simulations using a reduced number of individuals and SNPs (*N* = 8430 individuals and *M* = 14821 array SNPs; see Methods). We find that the single-component REML methods (GREML, BOLT-REML, and LDAK) are sensitive to MAF- and LD-dependent architectures that deviate from their respective model assumptions, whereas our estimator is robust to all architecture parameters. For example, when 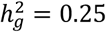 (Figure 3), GREML and BOLT-REML are accurate under the “single-component GREML model” (GREML: relative bias = −1.4%, *p* = 6.0 × 10^−3^, Supplementary Table S18; BOLT-REML: relative bias = −0.16%, *p* = 0.75, Supplementary Table S19) and LDAK is approximately unbiased under the “LDAK model” (relative bias = 0.16%, *p* = 0.77, Supplementary Table S20), but all single-component methods are sensitive to the MAF spectrum of causal variants and to the coupling of causal effects with MAF/LD. Across the 12 architectures in Figure 3 where *p*_causal_ = 1%, the relative biases of the single-component methods range from −15% to 7.9% (GREML), −14% to 9.1% (BOLT-REML), and −34% to 8.2% (LDAK) (Supplementary Tables S18-S20). In contrast, for the same 12 architectures, 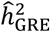 yields relative biases in the range [−2.1%, 1.7%], which is comparable to the relative bias observed with GREML-LDMS-I (range [−2.9%, 1.5%]) when using 8 GRMs defined by 4 LD quartiles and 2 MAF bins (MAF > 5% and MAF ≤ 5%) that align with the causal variant MAF spectrum (Figure 3, Supplementary Tables S21, S22). These trends are consistent across a range of values of 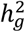 and *p*_causal_: across the 112 distinct LDAK-LD- and/or MAF-dependent architectures shown in Supplementary Figures S15-S19, the average and range of the relative bias are 0.09% [−4.9%, 6.4%] (GRE), −0.6% [−5.9%, 2.3%] (GREML-LDMS-I), −2.9% [−27%, 15%] (GREML), −1.8% [−25%, 18%] (BOLT-REML), and −8.2% [−44%, 13%] (LDAK) (Supplementary Tables S18-S22). Similar trends are observed in additional simulations under 14 LD-score-dependent architectures (Supplementary Figure S20, Supplementary Table S23). We note that the performance of GREML-LDMS-I depends on the resolution of the MAF and LD bins used to partition SNPs; in an extreme example where all causal variants are drawn from a MAF range tightly concentrated near 1%, running GREML-LDMS-I with the same 8 GRMs as before yields downward-biased estimates whereas our estimator remains robust (Supplementary Figure S21, Supplementary Tables S18-S22). While the variance of our estimator is larger than the variances of the REML-based methods (Figure 3), our approach is designed for biobank-scale GWAS data with sample sizes several orders of magnitude larger than what we used in these small-scale simulations. In summary, our results confirm that it is possible to accurately estimate 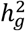 under minimal assumptions about genetic architecture.

**Figure 3.**
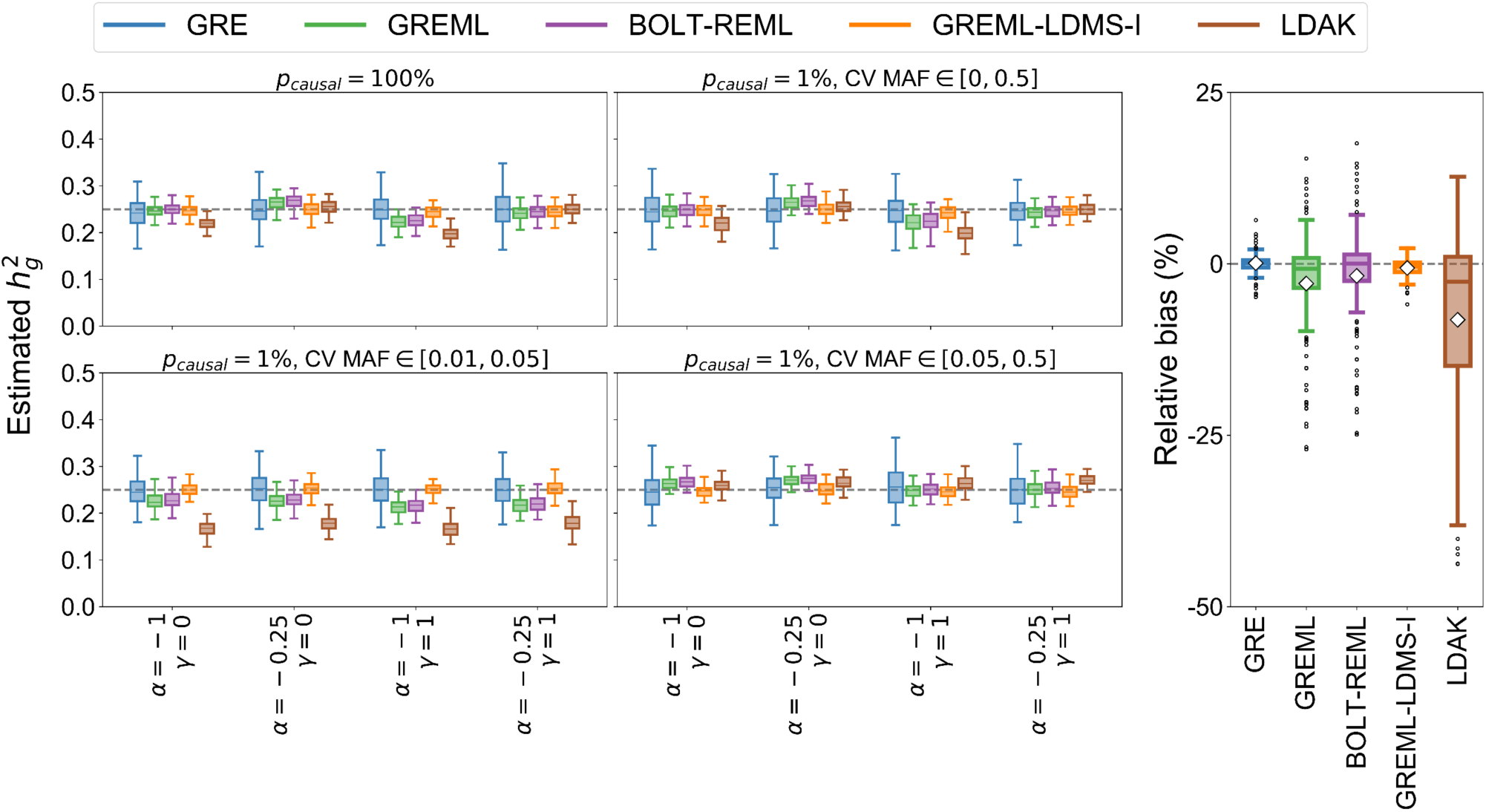
Comparison of 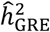 with GREML, BOLT-REML, GREML-LDMS-I, and LDAK in small-scale simulations (*N* = 8430 unrelated individuals, *M* = 14821 array SNPs). **Left:** Phenotypes were drawn under one of 16 MAF- and/or LDAK-LD-dependent architectures by varying *p*_causal_, α, *γ*, and CV MAF (see Methods). Each boxplot contains estimates of 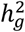 from 100 simulations. Boxplot whiskers extend to the minimum and maximum estimates located within 1.5 × IQR from the first and third quartiles, respectively. **Right:** Relative bias of each method (as a percentage of the true 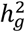) across 112 distinct MAF- and LDAK-LD-dependent architectures (see Methods). Each box plot represents the distribution of 112 points; each point is the average estimated 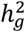 from 100 simulations under a single genetic architecture. The white diamonds mark the average of each distribution.

### Estimating SNP-heritability of 22 complex traits in the UK Biobank

Finally, we apply our approach to estimate 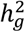 for 22 complex traits and diseases in the UK Biobank (*N* = 290K unrelated British individuals, *M* = 460K array SNPs; see Methods)^10^. For comparison, we also provide estimates of 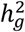 from LDSC (no annotations), S-LDSC (controlling for the baseline-LD model^13,30^), and SumHer. Of the 22 traits analyzed (6 quantitative and 16 binary), we focus on 18 traits for which 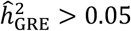 (Table 2). Using our approach, estimates of SNP-heritability for the 6 quantitative traits range from 0.12 (smoking status) to 0.60 (height). Across the 12 binary traits, our estimates range from 0.064 (autoimmune disorders) to 0.16 (hypertension) (Table 2). These estimates are robust to filtering of individuals based on relatedness (Supplementary Table S24), suggesting that including the top 20 PCs as covariates in OLS sufficiently controls for substructure and/or cryptic relatedness. We also computed 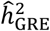 from two additional sets of SNPs (MAF > 0.1% and MAF > 0.01%) and found that the estimates increase slightly for lower MAF thresholds (Supplementary Table S25), which is expected due to the increased number of SNPs (the limited number of typed SNPs in the UK Biobank prohibits us from assessing the utility of GRE at rare variants further). To enable a direct comparison between 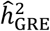 and the SNP-heritability quantities estimated by LDSC, S-LDSC, and SumHer, we run each of the summary-statistics-based methods with LD scores and regression weights computed from in-sample LD among the typed SNPs, and we estimate SNP-heritability as the sum of the per-SNP variances across all *M* SNPs (Methods). Across the 18 traits, the median difference (as a percentage of 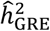) between S-LDSC (baseline-LD/in-sample) and 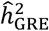 is −9%; the median difference between SumHer (in-sample) and 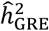 is 11% (Figure 4, Table 2). This pattern is roughly consistent with the global trends we observed in genome-wide simulations (Figure 2). As expected^11^, LDSC (in-sample) yields inflated estimates across all 18 traits.

**Table 2.** Estimates of 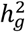 from the GRE approach, LDSC (in-sample), S-LDSC (baseline-LD/in-sample), and SumHer (in-sample) for 22 complex traits and diseases in the UK Biobank (*N* = 290K unrelated British individuals, *M* = 460K typed SNPs).

| Trait | GRE | S.E. | LDSC | S.E. | S-LDSC | S.E. | SumHer | S.E. |
| --- | --- | --- | --- | --- | --- | --- | --- | --- |
| Smoking Status | 0.122 | 3.90E-03 | 0.178 | 7.70E-03 | 0.110 | 8.50E-03 | 0.132 | 4.30E-03 |
| Height | 0.602 | 4.70E-03 | 0.730 | 2.70E-02 | 0.555 | 3.10E-02 | 0.634 | 2.70E-02 |
| BMI | 0.285 | 4.20E-03 | 0.436 | 1.20E-02 | 0.289 | 1.70E-02 | 0.315 | 9.00E-03 |
| WHR | 0.173 | 4.00E-03 | 0.256 | 1.20E-02 | 0.184 | 1.60E-02 | 0.198 | 9.40E-03 |
| Systolic Blood Pressure | 0.159 | 4.20E-03 | 0.243 | 9.00E-03 | 0.134 | 9.70E-03 | 0.177 | 5.70E-03 |
| Diastolic Blood Pressure | 0.154 | 4.20E-03 | 0.233 | 8.60E-03 | 0.130 | 9.70E-03 | 0.170 | 6.40E-03 |
| Eczema | 0.116 | 4.20E-03 | 0.165 | 1.10E-02 | 0.107 | 1.20E-02 | 0.130 | 8.80E-03 |
| Asthma | 0.116 | 4.90E-03 | 0.163 | 1.20E-02 | 0.116 | 1.70E-02 | 0.131 | 1.20E-02 |
| Hypertension | 0.162 | 4.00E-03 | 0.244 | 9.40E-03 | 0.142 | 1.10E-02 | 0.180 | 6.10E-03 |
| High Cholesterol | 0.082 | 5.10E-03 | 0.127 | 1.30E-02 | 0.138 | 5.80E-02 | 0.088 | 8.30E-03 |
| Diabetes (Any) | 0.070 | 3.70E-03 | 0.093 | 5.90E-03 | 0.062 | 8.70E-03 | 0.074 | 5.00E-03 |
| Type 2 Diabetes | 0.071 | 3.80E-03 | 0.090 | 6.10E-03 | 0.057 | 8.80E-03 | 0.071 | 4.00E-03 |
| Hypothyroidism | 0.088 | 5.20E-03 | 0.142 | 1.30E-02 | 0.078 | 1.20E-02 | 0.110 | 1.70E-02 |
| Thyroid Disorders | 0.084 | 5.20E-03 | 0.141 | 1.30E-02 | 0.080 | 1.20E-02 | 0.110 | 2.00E-02 |
| Endocrinopathies | 0.069 | 5.10E-03 | 0.084 | 7.00E-03 | 0.058 | 9.90E-03 | 0.068 | 5.00E-03 |
| Cardiovascular Diseases | 0.143 | 5.30E-03 | 0.228 | 1.10E-02 | 0.140 | 1.40E-02 | 0.164 | 6.00E-03 |
| Respiratory and ENT Diseases | 0.086 | 5.20E-03 | 0.120 | 1.20E-02 | 0.079 | 1.40E-02 | 0.090 | 9.50E-03 |
| Psoriasis | 0.019 | 5.00E-03 | 0.071 | 3.10E-02 | 0.035 | 1.20E-02 | 0.059 | 4.20E-02 |
| Dermatologic Disorders | 0.023 | 5.00E-03 | 0.049 | 1.40E-02 | 0.034 | 9.90E-03 | 0.031 | 1.10E-02 |
| Rheumatoid Arthritis | 0.008 | 5.00E-03 | 0.041 | 2.10E-02 | 0.010 | 7.90E-03 | 0.021 | 1.20E-02 |
| Autoimmune Disorders (Broad) | 0.063 | 5.10E-03 | 0.105 | 1.20E-02 | 0.050 | 9.50E-03 | 0.079 | 1.70E-02 |
| Autoimmune Disorders (Certain) | 0.015 | 5.00E-03 | 0.052 | 2.60E-02 | 0.005 | 7.60E-03 | 0.047 | 3.40E-02 |

**Figure 4.**
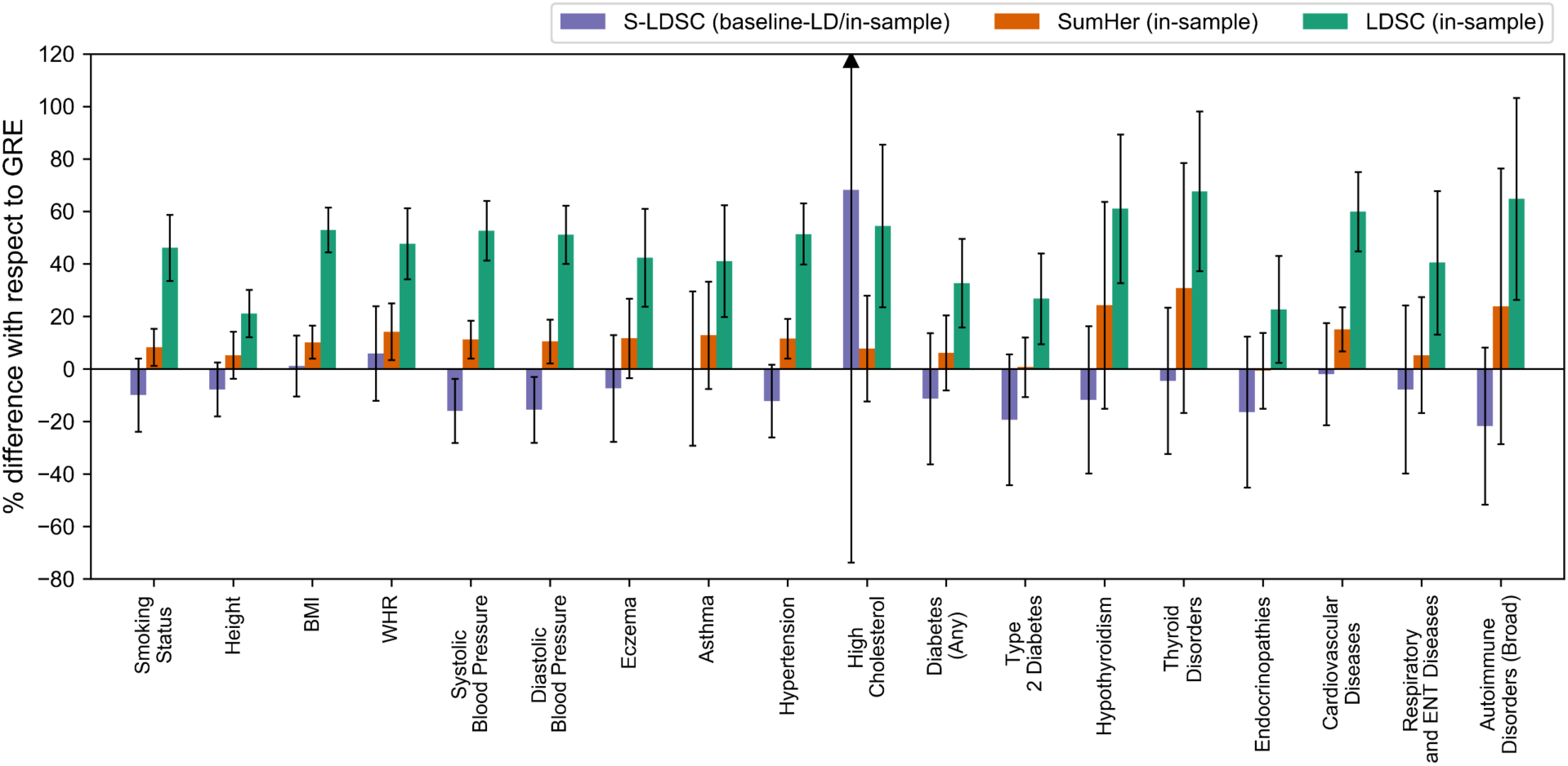
Percent difference of 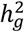 estimates from LDSC (in-sample), S-LDSC (baseline-LD/in-sample), and SumHer (in-sample) with respect to 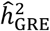 for 18 complex traits and diseases in the UK Biobank for which 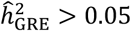 (*N* = 290K unrelated British individuals, *M* = 460K typed SNPs; see Methods). Each bar represents the difference between the estimated 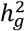 from one of the methods (LDSC, S-LDSC, or SumHer) and 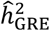 as a percentage of 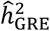. Black bars mark ±2 standard errors.

To enable a comparison between 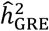 and SNP-heritability estimates from summary-statistics-based methods reported in the literature, we also run LDSC, S-LDSC, and SumHer with their recommended parameter settings^11,12,14,30^ and with LD scores and regression weights computed from 489 Europeans in the 1000 Genomes Phase 3 reference panel^31^ – we note that when running these methods as recommended, their SNP-heritability estimands are not equivalent to our definition of 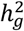 (see Methods and refs.^11,12,14,19^ for details). Across the 18 traits for which 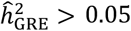, the median differences with respect to 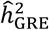 are −11% for LDSC (1KG), −14% for S-LDSC (baseline-LD/1KG), and 38% for SumHer (1KG) (Supplementary Figure S22, Supplementary Table S26). Across 9 traits (a subset of the 18 traits with 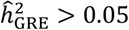) for which a previous study reported estimates from single-component BOLT-REML (computed from approximately 337K unrelated white British individuals in the UK Biobank^27^), the median difference between the previously reported BOLT-REML estimates and 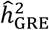 is 8% (Supplementary Table S26).

### Runtime and memory requirement

Since our approach is designed to be applied to biobank-scale data, we report the runtime and memory requirements for computing 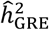 with 22 chromosome-wide LD blocks in the UK Biobank (*N* = 337K individuals, *M* = 593K array SNPs). First, we compute chromosome-wide LD; this has complexity 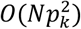 for chromosome *k* with *p*_*k*_ SNPs. In practice, this step does not impose a computational bottleneck because the LD computations can be parallelized over SNP partitions (e.g., for a pair of SNP partitions containing 1000 SNPs each, computing the pairwise LD matrix takes about 10 minutes and 16GB of memory). Second, the pseudoinverse of each chromosome-wide LD matrix is computed via truncated singular value decomposition (SVD), which has complexity 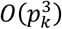 for chromosome *k*. This step is parallelized over chromosomes; for chromosome 2, which has the largest number of typed SNPs, computing the truncated SVD and pseudoinverse of the LD matrix takes about 3 hours and 60GB of memory. Lastly, given the precomputed pseudoinverse of each chromosome-wide LD matrix and OLS association statistics, computing 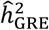 genome-wide has complexity 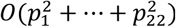. For any of the UK Biobank traits analyzed in this work, this takes less than 1 hour and requires 24GB of memory; most of this time is spent loading the pseudoinverse LD matrices into memory. For comparison, running LDSC, S-LDSC, or SumHer consists of precomputing LD scores and SNP-specific weights and performing linear regression to estimate the variance parameters in the model. The first step (precomputing LD scores and SNP-specific weights) can be parallelized over blocks of SNPs and therefore does not pose a computational bottleneck in practice. The complexity of the second step (least squares regression) is *O*(*C*^2^*M*) where *M* is the number of SNPs in the regression and *C* is the number of variance parameters being estimated.

## Discussion

In this work, we show that highly accurate estimation of SNP-heritability can be achieved under minimal assumptions on the genetic architecture of complex traits. In particular, our proposed estimator assumes that each SNP effect has a fixed SNP-specific variance that can capture any arbitrary relationship between effect size and genomic features such as MAF and LD. We show that all existing methods to estimate SNP-heritability impose additional assumptions on the GRE model, and we confirm through extensive simulations that these methods are susceptible to bias when their modeling assumptions are not met. Additionally, we confirm that REML-based methods that partition SNPs by MAF and LD score generally yield much smaller bias compared to single-component REML methods^18^. In contrast, our estimator, derived under the GRE model, provides accurate estimates of SNP-heritability regardless of the underlying genetic architecture, without specifying a heritability model or partitioning SNPs by functional categories. On average across 18 heritable traits in the UK Biobank 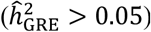, our approach yields estimates that are higher than S-LDSC estimates (controlling for the baseline-LD model^30^) and lower than SumHer estimates (with the recommended heritability model^9,14^). One practical advantage of our approach over methods such as LDSC, S-LDSC, and SumHer is that the estimand of our approach is always the same for a given genotype matrix, whereas the definitions and interpretations of the estimands of LDSC, S-LDSC, and SumHer can vary depending on what sets of SNPs are used in each step of the inference procedure (e.g., the set of SNPs used to compute LD scores need not be the same set of SNPs that defines the SNP-heritability estimand of interest)^11,12,19^. Overall, our results show that while existing methods can yield biases, for the purpose of estimating total SNP-heritability of complex traits, most methods are relatively accurate and robust to plausible genetic architectures.

We conclude with several caveats and future directions. First, the utility of the GRE estimator critically depends on the ratio between the number of SNPs (*M*) and the number of individuals (*N*) in the data – as *M*/*N* increases, the eigenstructure of the in-sample LD matrix (and sample covariance matrices in general) becomes increasingly distorted (larger eigenvalues are overestimated and smaller eigenvalues are underestimated)^32^. We mitigate this by assuming that the genome-wide LD matrix has a block diagonal structure (specifically, one block per chromosome); since the number of unrelated British individuals in the UK Biobank is larger than the number of array SNPs per chromosome, our approach is able to provide meaningful estimates of the SNP-heritability attributable to common SNPs (MAF > 1%) in individuals of British ancestry. While the utility of our approach is clearly limited for the time being by the availability of individual-level biobank-scale data, this will become less of a concern as more institutions establish their own biobanks^33–35^. A major limitation of our approach remains with respect to imputed and/or whole-genome sequenced data, in which the number of SNPs will continue to be orders of magnitude larger than the number of individuals for the foreseeable future. We defer a thorough investigation of regularized estimation of LD in high-dimensional settings (*M > N*) to future work.

Second, the theoretical guarantees of the GRE estimator rely on the assumption that OLS association statistics and chromosome-wide LD matrices are estimated from the same genotype data. While summary statistics have been made publicly available for hundreds of large-scale GWAS, in-sample LD is usually unavailable or nonexistent for these studies since most are meta-analyses^36^, and publicly available reference panels such as 1000 Genomes^31^ currently have sample sizes in the hundreds or thousands at most. In addition, many publicly available summary statistics were computed using linear mixed models rather than OLS in order to control for population structure/cryptic relatedness. Previous works have noted (in the context of statistical fine-mapping) that the LD computation must be adjusted to accommodate association statistics computed from mixed models^36,37^. The sensitivity of our estimator to reference panel LD (with or without regularized LD estimation) and/or mixed model association statistics remains unclear^29,38^; we leave an investigation of both for future work. Furthermore, our simulations use typed SNPs to draw phenotypes because imputed genotypes have highly irregular LD patterns^9,18^. Although it would be more realistic to simulate causal variants and phenotypes from a denser set of genotyped SNPs or from whole-genome sequencing data^18^, our simulation design was dependent on the availability of individual-level genotype measurements in biobank-scale sample sizes.

Third, the GRE estimator does not correct for population structure/cryptic relatedness. We mitigate this in our analysis of real UK Biobank traits by considering only unrelated individuals (> 3rd degree relatives) and by including age, sex, and the top 20 principal components as covariates in the linear regression when computing OLS association statistics. While recent work has found significant evidence of assortative mating for some traits in the UK Biobank (e.g., height) and not others^39^, our estimates for real phenotypes are robust to different relatedness thresholds, suggesting that including the top 20 PCs as covariates in OLS is sufficient to control for population stratification. Still, it remains unclear how to quantify the bias of our genome-wide estimator due to population structure and/or assortative mating in real data. In addition, we derive the GRE estimator under no ascertainment in case/control data. Future work is needed to extend the GRE approach to control for ascertainment bias^15,16,40,41^.

Finally, while previous works have applied similar estimators in the context of fixed effects models to estimate local SNP-heritability within small regions (e.g., LD blocks)^28,29^, additional work is needed to extend our approach to perform functional partitioning of SNP-heritability by higher-resolution annotations. Existing methods for partitioning genome-wide SNP-heritability by small and/or overlapping annotations make various assumptions on genetic architecture^8,12–14,30^, motivating the development of new methods in this area under fewer assumptions.

## URLs

GRE estimator: https://bogdan.dgsom.ucla.edu/pages/software

BOLT-LMM: https://data.broadinstitute.org/alkesgroup/BOLT-LMM/

GCTA: https://cnsgenomics.com/software/gcta/

LDAK: http://dougspeed.com/ldak/

LDSC: https://github.com/bulik/ldsc/

baseline-LD annotations: https://data.broadinstitute.org/alkesgroup/LDSCORE/

PLINK: https://www.cog-genomics.org/plink2

UK Biobank: https://www.ukbiobank.ac.uk

## Acknowledgments

This research was conducted using the UK Biobank Resource under applications 33297 and 33127. We thank the participants of UK Biobank for making this work possible. We also thank Ruth Johnson, Malika Kumar Freund, Megan Major, Steven Gazal, Alkes Price and David Balding for helpful discussions. This work was funded by the National Institutes of Health (NIH) under awards R01HG009120, R01MH115676, R01HG006399, U01CA194393, T32NS048004, T32MH073526, and T32HG002536.

## Methods

### The generalized random effects model

We model the phenotype for an individual *n* randomly sampled from the population as 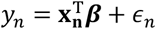, where **x_n_** = (*x*_*n*1_ … *x*_*nM*_) ^*T*^ is a vector of standardized genotypes measured at *M* SNPs for individual *n*, ***β*** = (*β*_1_, …, *β*_*M*_)^*T*^ is an *M*-vector of the corresponding standardized SNP effect sizes, and 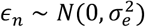 is environmental noise. We assume Var[*y*_*n*_] = 1 and that the genotype at each SNP *i* is centered and scaled in the population such that E[*x*_*ni*_] = 0 and Var[*x*_*ni*_] = 1; i.e. 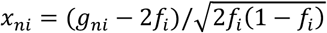, where *g*_*ni*_ ∈ {0,1,2} is the number of copies of the effect allele at SNP *i* for individual *n*, and *f*_*i*_ is the population frequency of the effect allele at SNP *i*. We define the population LD between two SNPs *i* and *j* to be *ν*_*ij*_ ≡ E[*x*_*ni*_*x*_*nj*_] for all *i* ≠ *j*. The population LD matrix among the *M* SNPs is therefore 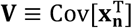. For simplicity, we use “SNP effect sizes” in lieu of “standardized SNP effect sizes” to refer to ***β***. We assume that the genotypes **x_n_** and effect sizes ***β*** are independent given allele frequencies (*f*_1_, …, *f*_*M*_) and **V**.

Under the generalized random effects (GRE) model, the first two moments of the distribution of the effect size of SNP *i* are E[*β*_*i*_] = 0 and 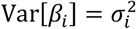, where 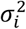 can be any arbitrary nonnegative finite number. We assume the covariance between the effects of different SNPs is 0 (i.e. Cov;*β*_*i*_, *β*_*j*_ = = E;[*β*_*i*_*β*_*j*_]= = 0 for all i ≠ j). Because the SNP-specific variances 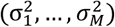 can capture any polygenicity (number of variants with effects larger than some measurable constant) and any degree of coupling between genomic features (e.g., MAF and LD) and effect size, the GRE model encompasses most realistic genetic architectures (Table 1).

We define total SNP-heritability 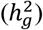 to be the proportion of phenotypic variance attributable to the additive effects of a set of *M* SNPs whose genotypes are directly measured:

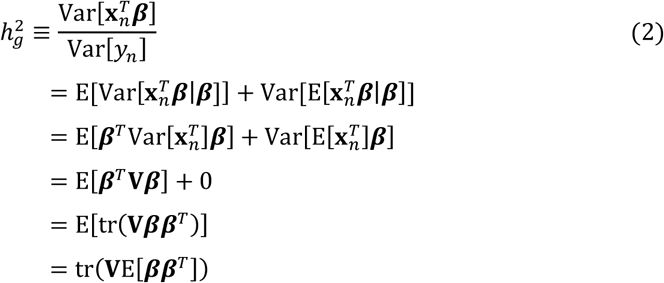

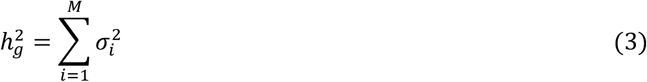

Thus, 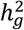 is defined with respect to a given population and a given set of SNPs. By definition, 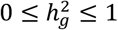. Similarly, we define regional SNP-heritability 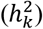 to be the proportion of phenotypic variance due to the additive effects of the genotyped SNPs in region *k*. We assume that the set of SNPs that defines 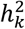 is a subset of the *M* SNPs that define 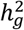 (thus, 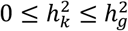). If region *k* is the whole genome, 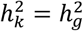.

### Estimating SNP-heritability under the GRE model

We are interested in estimating 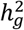 under the GRE model (Equation 3). In a GWAS with *N* individuals genotyped at *M* SNPs, let 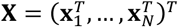 be the *N* × *M* matrix of standardized genotypes (i.e. each column of ***X*** has been standardized to have mean 0 and variance 1), let ***y*** *=* (*y*_1_, …, *y*_*N*_)^*T*^ be the *N*-vector of standardized phenotypes, and let 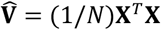 be the *M* × *M* in-sample LD matrix (an estimate of population LD, ***V***) with rank *q*, where 1 ≤ *q* ≤ *M*. Let ***X*** *=* (***X***_1_, …, ***X***_*K*_) be the genotype matrices for a set of *K* approximately independent regions spanning all *M* SNPs (e.g., chromosomes). For each region *k* containing *p*_*k*_ SNPs, ***X***_*k*_ is the *N* × *p*_*k*_ standardized genotype matrix and 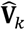 is the corresponding *p*_*k*_ × *p*_*k*_ in-sample LD matrix with rank *q*_*k*_, where 1 ≤ *q*_*k*_ ≤ *p*_*k*_. We propose the following estimator for genome-wide SNP-heritability:

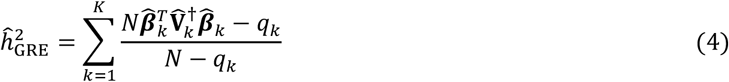

where 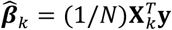 is the *p*_*k*_-vector of marginal SNP effects estimated by ordinary least squares (OLS) for region *k* and 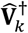 is the pseudoinverse of 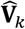.

In the following sections, we first derive 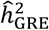 in the simplest case where *K =* 1 and *N* > *M* by finding an estimator that satisfies 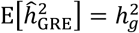. We then describe modifications to this estimator to allow *N* < *M* as well as rank-deficient LD matrices. Lastly, we derive an analytical form for the standard error of 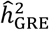.

### Derivation for 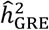 assuming fixed *β* and *N* > *M*

Recall that Var[*y*_*n*_] *=* 1 and 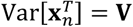. Our goal is to find an estimator 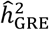 that satisfies 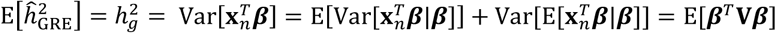 (Equation 2). If ***β*** were fixed and we observed ***V*** and ***β***, we could estimate 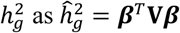. However, in reality, we observe noisy estimates of ***β*** and ***V*** from GWAS. Given a GWAS of *N* unrelated individuals and*M* SNPs, we observe ***X***, the standardized genotype matrix, and ***y***, the standardized phenotype vector. We assume that when 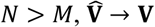 as *N* →∞ (in practice, the assumption that *N* > *M* is untrue; in subsequent sections we show how we partition the genome into*K* blocks such that *N* > *p*_*k*_ for each block *k*). In a typical GWAS, the marginal SNP effects are estimated through ordinary least squares (OLS) regression as. 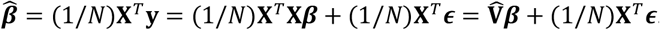. Given ***X*** and fixed ***β***, it follows that

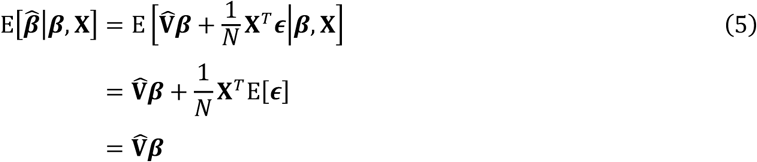

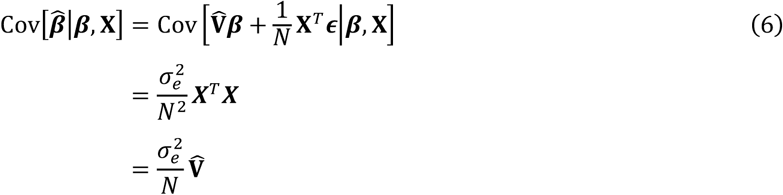

Thus, as 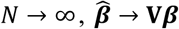 substituting 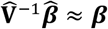 and 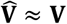, we obtain the revised estimator 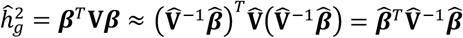. The expectation of this estimator is

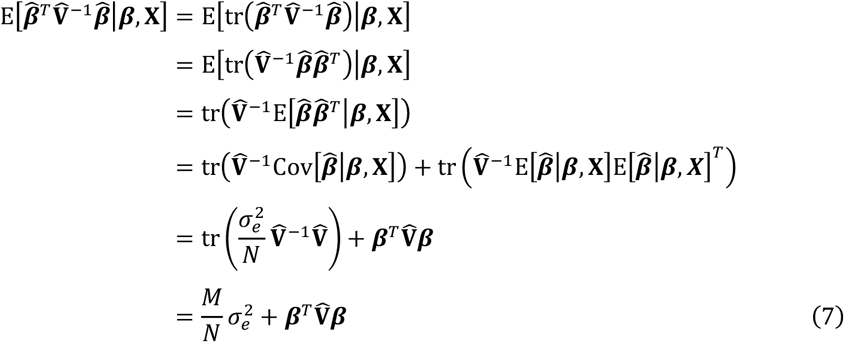

We define 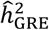 to be an estimator that satisfies 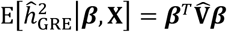. Substituting into Equation 7, we obtain

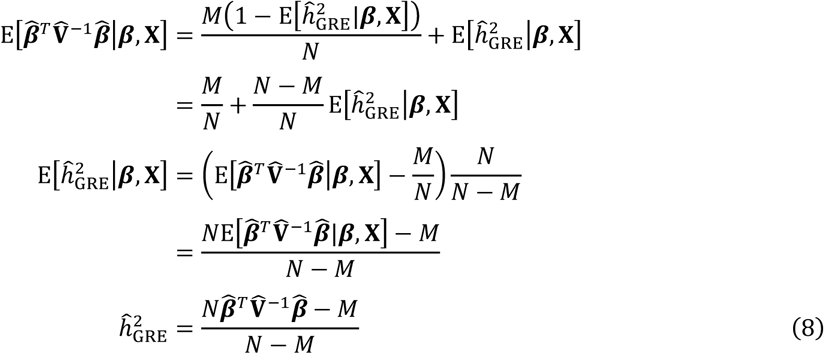

### Unbiasedness of 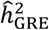 under the GRE model when *N* > *M*

Recall that under the GRE model, E[*β*_*i*_] *=* 0 and 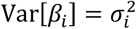, where 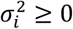 for all SNPs *i*. In the previous sections, we showed that 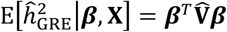 and 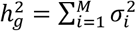. Recalling that *cov*[*β*_*i*_ *β*_*j*_]=0 for all *i* ≠ *j*, it follows that

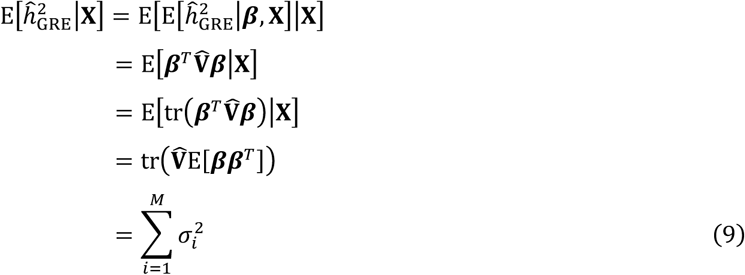

Therefore, 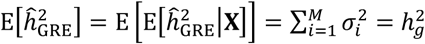. This implies that 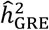 is an unbiased estimator for 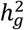 under a wide range of genetic architectures that fall under the GRE model.

### Genome-wide approximation

For most GWAS, because the number of genotyped SNPs *M* is much larger than the number of individuals *N* in the study, 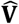 is a poor estimator of ***V*** genome-wide; as *M*/*N* increases, the eigenstructure of 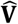 becomes increasingly distorted (larger eigenvalues are overestimated and smaller eigenvalues are underestimated)^32^. In addition, it is generally computationally intractable to compute and invert 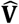 genome-wide. Thus, in practice, we divide the genome into a set of *K* approximately independent blocks (e.g., by chromosome) and, following a procedure similar to Equations 5-8, we obtain

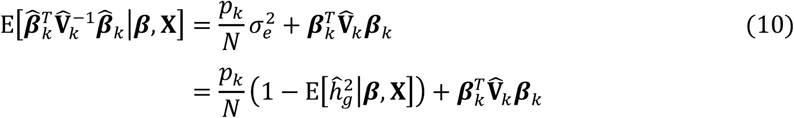

To find an estimator that satisfies 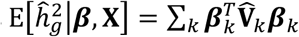, we sum Equation 10 over *k =* 1, …, *K*:

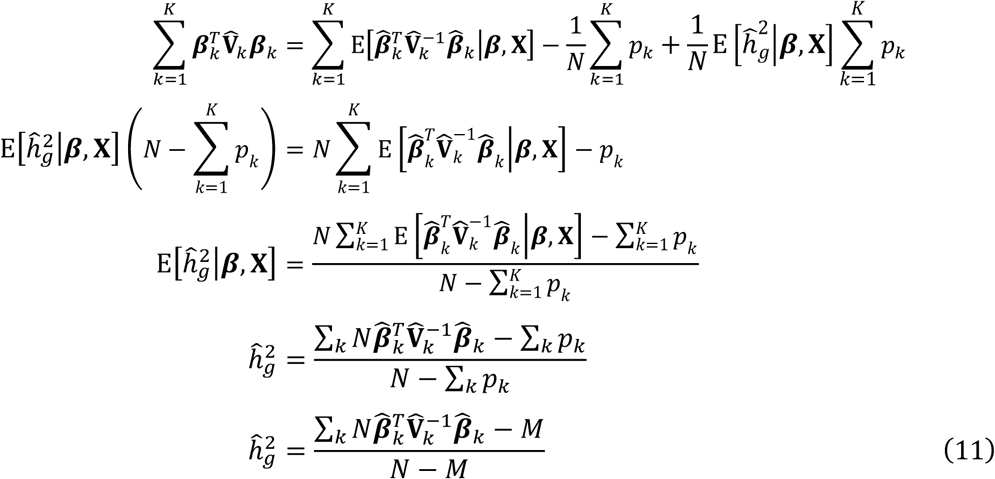

While Equation 11 does circumvent the need to invert the genome-wide LD matrix in Equation 8, this estimator will produce negative estimates of 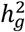 if *N* < *M*, which is the case in all of our genome-wide analyses. We therefore use an approximation which estimates the contribution of block *k* while ignoring the contributions of the remaining blocks. That is, assuming ***y*** *=* ***X***_*k*_***β***_*k*_ + **ϵ**_*k*_, where 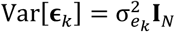, we obtain

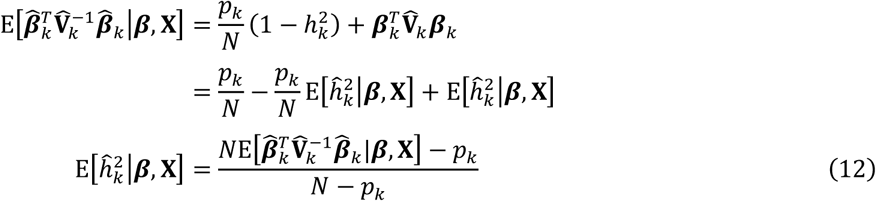

An estimator that satisfies Equation 12 is

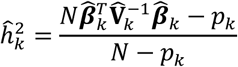

Finally, we estimate genome-wide SNP-heritability as

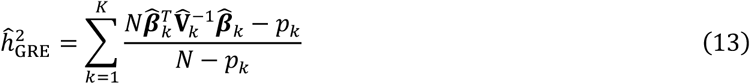

While this estimator does not provide theoretical guarantees of unbiasedness, we find that it allows us to robustly estimate genome-wide SNP-heritability as long as *N* ≫ *p*_*k*_ for all *k* (e.g., Figure 1b).

### Extension for rank-deficient LD

It is often the case that two SNPs are perfectly correlated in a genotype block ***X***_*k*_, or that *N* < *p*_*k*_ for a block*k*. In this case, 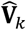 is rank-deficient (i.e. its rank is less than *p*_*k*_) and 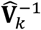 does not exist. We therefore compute 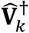, the pseudoinverse (Moore-Penrose inverse) of 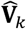, which approximate 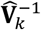 using its truncated eigendecomposition. Let 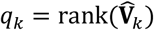 and let 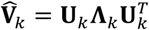 be the eigendecomposition of 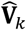, where 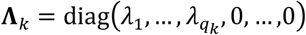. The pseudoinverse of 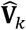 is 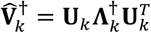, where 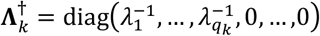.

Substituting 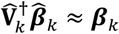 and 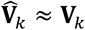, we obtain the following estimator for 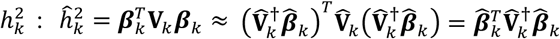. Let 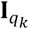 be a *p*_*k*_ × *p*_*k*_ diagonal matrix in which the first *q*_*k*_ diagonal entries are 1 and the rest are 0. The expectation of our estimator given ***β*** and ***X*** is

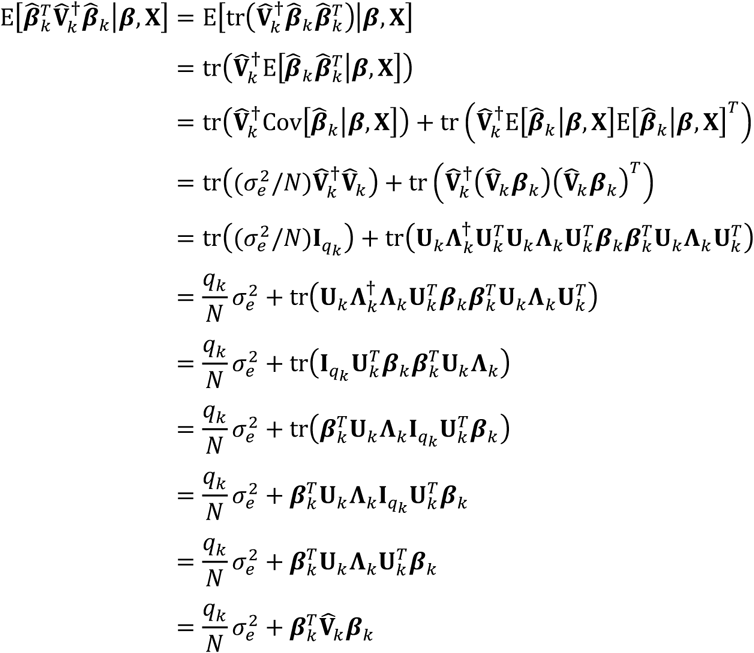

Following a procedure similar to Equations 10-13, we obtain

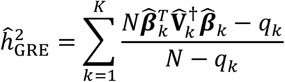

Again, while this estimator is not unbiased, it allows us to robustly estimate genome-wide SNP-heritability as long as *N* ≫ *q*_*k*_ for all *k*.

### Analytical variance of 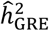

Following quadratic form theory^29,42^, the variance of 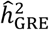 in the single-block case is given by

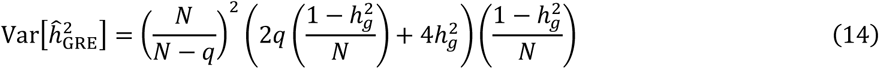

When using the *K*-block approximation, which assumes that the blocks are independent, we approximate Equation 14 as the sum of the variances of the local SNP-heritabilities:

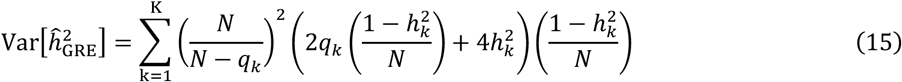

Because 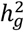 and 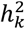 for all *k* are unknown, Equation 14 is estimated by plugging in 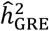 and Equation 15 is estimated by plugging in 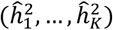, the estimates of the regional SNP-heritabilities.

### Simulation Framework

To assess the performance of 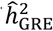 and other methods, we simulated continuous phenotypes from genotype array data in the UK Biobank^10^ under a range of genetic architectures. We obtained a set of *N* = 337205 unrelated British individuals to use in simulations by extracting individuals that are > 3rd degree relatives (defined as pairs of individuals with kinship coefficient < 1/2^(^9^/^2^)^)^10^ and excluding individuals with putative sex chromosome aneuploidy. In all simulations, we standardize the genotype matrix before drawing phenotypes such that each column (SNP) of the genotype matrix has mean 0 and variance 1. In other words, we standardize the genotype at SNP *i* for individual *n* by computing 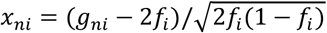, where *g*_*ni*_ *∈* {0,1,2} is the number of minor alleles at SNP *i* for individual *n* and *f*_*i*_ is the minor allele frequency (MAF) of SNP *i* among the *N* individuals.

#### Simulations of quantitative traits with no population stratification

Given standardized genotypes for *N* individuals at *M* SNPs and a fixed value of 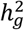 phenotypes are simulated under different genetic architectures according to the following model. The proportion of causal variants, *p*_causal_, is set to either 1 (i.e. an infinitesmal model in which all variants have nonzero effects), 0.01, or 0.001. Let *c*_*i*_ ∈ {0,1} be an indicator variable for the causal status of SNP *i*. If *p*_causal_ = 1, *c*_*i*_ = 1 for *i* = 1, …, *M*. Otherwise, if 0 *≤ p*_causal_ < 1, we draw *p*_causal_ *× M* SNPs from the set of SNPs with minor allele frequencies in one of three ranges: (0, 0.5], (0.01, 0.05], or (0.05, 0.5]. We use the abbreviation “CV MAF” to refer to the MAF range from which causal variants are drawn. The standardized SNP effect sizes and phenotypes are then drawn according to the following model:

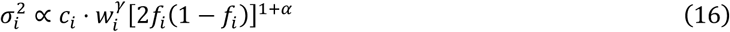

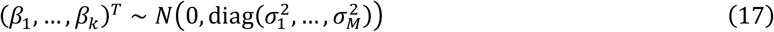

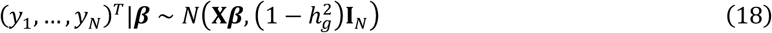

where *α* is a parameter that controls the coupling of MAF and effect size, *w*_*i*_ is a SNP-specific LD weight, and *γ* ∈ {0,1}is a global parameter specifying whether the effect size of a SNP is coupled with its LD score. We simulate two types of LD-dependent architectures by defining the SNP-specific LD weights *w*_1_, …, *w*_*M*_ to be either (1) the default “LDAK weights” computed by the LDAK software^6^, or (2) the inverse unpartitioned “LD score” of each SNP computed within a 2-Mb window using the LDSC software (i.e. 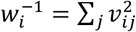 here *j* indexes the set of SNPs within a 2-Mb window centered on SNP *i*)^11^. When γ = 1, both the LDAK weights and inverse LD score weights cause SNPs in regions of higher LD to have smaller effects than do SNPs in regions of lower LD. We set α to one of two values: α = −1, which indicates a relatively strong inverse relationship between MAF and effect size, or α = −0.25, which indicates a weaker inverse relationship between MAF and effect size. Each per-SNP variance is multiplied by a constant scaling factor to ensure that 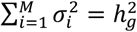. Note that 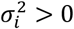 if *c*_*i*_ = 1 and 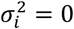 if *c*_*i*_ = 0.

Finally, given simulated phenotypes y = (*y*_1_, …, *y*_*N*_)^*T*^and genotypes 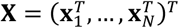, we compute marginal association statistics through ordinary least squares (OLS) as 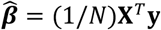.

#### Simulations of case-control phenotypes with no population stratification

To simulate case-control studies, we first draw each individual’s continuous liability (*l*_*n*_ for individual *n*) according to Equation 18. Then, for a given population prevalence (0 *≤ d*_*pop*_ *≤* 1), we compute the corresponding liability threshold *L* = Φ^−1^(1 *– d*_*pop*_), where Φ is the CDF of the standard normal distribution, and we convert each individual’s continuous liability into a case-control status: *y*_*n*_ = 1 if *l*_*n*_ *≥ L* or *y*_*n*_ = 0 if *l*_*n*_ < *L*. In simulations of unascertained case-control studies, we assume that the proportion of cases in the study is equal to the population prevalence (*d*_*GWAS*_ = *d*_*pop*_). In all simulations of ascertained case-control studies (*d*_*GWAS*_ *> d*_*pop*_), we set *d*_*GWAS*_ = 0.5 and select a random set of controls to satisfy *N*_*case*_= *N*_*control*._

To estimate SNP-heritability from simulated case-control studies, we compute association statistics by regressing the binary case-control statuses on genotypes and apply GRE; this produces an estimate of SNP-heritability on the *observed* scale 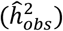. We assume that we know the population prevalence, which allows us to convert this fersotimmattehe observed scale to the *liability* scale with the transformation 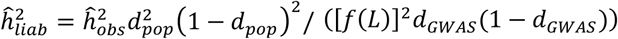, where *f* is the standard normal probability density function^43^.

#### Simulations with population stratification

To simulate GWAS with population stratification, we draw phenotypes from a model where a covariate that is correlated to genotypes has a nonzero effect on phenotype. To this end, we simulate an effect of the first genetic principal component (PC_1_) by setting 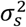, the proportion of total phenotypic variance explained by the covariate, and drawing phenotypes from the model

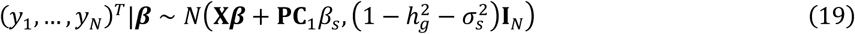

where *β*_*s*_ satisfies 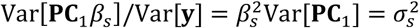. We then compute association statistics from one of two models: y = X^*T*^ *β +* ϵ, which ignores population stratification and other potential sources of confounding, or y = X^*T*^ *β +* PC_1_*β*_*s*_ *+* ϵ, which controls for the effect of the first genetic PC.

### Comparison of methods in simulations

Unless otherwise specified, in all genome-wide simulations, we use real genotypes of *N* = 337205 unrelated British individuals measured at *M* = 593300 array SNPs to draw causal effects for all *M* SNPs and phenotypes for all *N* individuals. OLS summary statistics are computed for all *M* SNPs using the simulated phenotypes and real genotypes for all *N* individuals. We implement our estimator (Equation 4) by computing chromosome-wide in-sample LD for each chromosome *k* as 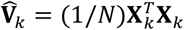 and we compare to three computationally efficient methods that operate on summary statistics: LD score regression (LDSC)^11^, stratified LD score regression (S-LDSC)^12,13^, and SumHer^14^.

To run LDSC with no annotations, we use the LDSC software (see URLs) to compute the LD score of each SNP as a function of its LD to all other SNPs in a 2-Mb window centered on the SNP. The LD scores are computed from a random sample of 40K individuals to reduce the amount of memory required by the LDSC software. We run the regression with an unconstrained intercept, using all *M* SNPs as observations in the response variable, where each SNP in the regression is weighted to account for heteroscedasticity and correlations between association statistics at SNPs in LD^11^. 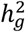 is estimated as a function of all *M* SNP-specific variances by running LDSC with the flags –not-M-5-50 and –chisq-max 99999 (the latter option prevents the LDSC software from dropping high-effect SNPs).

We run S-LDSC in two ways to account for MAF- and LD-dependent architectures. S-LDSC (MAF) refers to S-LDSC with 10 binary MAF bin annotations defined such that each bin contains exactly 10% of the typed SNPs; this is intended to mirror the 10 MAF bin annotations in the S-LDSC “baseline-LD model”^13^ (see Supplementary Table S14 for precise MAF bin ranges for the UK Biobank Axiom Array). S-LDSC (MAF+LLD) refers to S-LDSC with the same 10 MAF bins and an additional continuous “level of LD” (LLD) annotation computed by quantile-normalizing the unpartitioned LD scores within each MAF bin to a standard normal distribution^13^. While our definition of LLD is intended to mirror the LLD annotation in the baseline-LD model, we do not set the LLD of variants with MAF < 0.05 to 0 because our estimand of interest is the SNP-heritability attributable to all *M* SNPs (not just SNPs with MAF > 0.05)^13^. For each annotation, LD scores are computed within 2-Mb windows from a random sample of 40K individuals. We run the regression with all *M* SNPs, an unconstrained intercept, and the recommended regression weights^12,13^. Once again, we use the flags –not-M-5-50 and –chisq-max 99999 to estimate 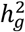 as a function of all *M* SNP-specific variances and to prevent the LDSC software from dropping high-effect SNPs.

To run SumHer, we first use the LDAK software (see URLs) to compute the default “LDAK weights” using in-sample LD^6,9,14^. Second, we compute “LD tagging” (i.e. LD scores) using 1-Mb windows centered on each SNP and setting α = −0.25 as recommended^14^. The LDAK software is memory-efficient, allowing us to use in-sample LD computed from all *N* = 337K individuals to obtain LDAK weights and LD tagging. Finally, we run SumHer to estimate 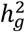 as a function of all *M* SNP-specific variances. Unless otherwise specified, all default parameter settings are used to run SumHer in simulations.

Similarly, in all small-scale simulations, we use real genotypes of *N* = 8430 unrelated individuals at *M* = 14821 array SNPs to draw phenotypes for all *N* individuals. These individuals and SNPs are a subset of the full UK Biobank data that were used in the genome-wide simulations, and were chosen by selecting approximately 2.5% of individuals and the first 2.5% of SNPs from the beginning of each chromosome in order to preserve a realistic LD structure among the SNPs. OLS summary statistics are computed from the simulated phenotypes and genotypes for all *N* individuals and *M* SNPs, and 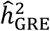 is computed using in-sample chromosome-wide LD. We run the implementation of single-component GREML^3^ provided by the GCTA software^44^ and single-component BOLT-REML^8^ provided by the BOLT-LMM software (see URLs), both with default parameters. We run the implementation of GREML-LDMS-I^18^ provided by the GCTA software using 8 GRMs created from 2 MAF bins (MAF *≤* 0.05 and MAF > 0.05) and 4 LD score quartiles; LD scores were computed using the GCTA software with the default window size of 200-kb. We run LDAK using the default LDAK weights, setting *α* = −0.25 as recommended^6,9^.

For a given genetic architecture, we generate 100 simulation replicates and obtain 100 estimates of *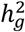* from each method. We estimate the bias of an estimator 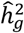 under a given architecture by computing the difference between the average of the 100 estimates and the simulated 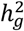 (i.e. bias 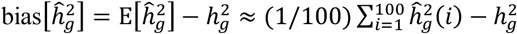 where 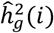 is the estimate from the *i*-th simulation). To test whether the bias is statistically significant (i.e. significantly different from 0), we assess the z-score of the 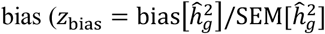, where 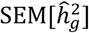 is the standard error of the mean of the 100 estimates) which follows a *N*(0,1) distribution under the null hypothesis. To enable a comparison of estimators across different values of 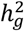, we assess the relative bias of an estimator under a single architecture 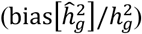 as a percentage of 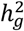. In Figure 1c, we compute the error of a single estimate from the *i*-th simulation as 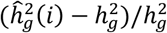 errors are also reported as percentages of 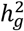.

We also performed simulations using the genotypes of 7,685 individuals of South Asian ancestry in the UK Biobank. This group was composed of individuals of Indian (*n* = 5,716), Pakistani (*n* = 1,748), and Bangladeshi (*n* = 221) ancestry. Due to the small sample size, we used a reduced set of 803 SNPs from chromosome 21 and 839 SNPs from chromosome 22 (1,642 SNPs in total). This reduced set of SNPs was chosen such that *N*/*p*_*k*_ for each chromosome *k* was similar to *N*/*p*_*k*_ in the “white British” cohort.

### Analysis of UK Biobank phenotypes

We estimate SNP-heritability for 22 real complex traits (6 quantitative, 16 binary) in the UK Biobank^10^. We use PLINK^45^ to exclude SNPs with MAF < 0.01 and genotype missingness > 0.01 as well as SNPs that fail the Hardy-Weinberg test at significance threshold 10^−7^. We keep only the individuals with self-reported British white ancestry and no kinship (i.e. > 3rd degree relatives, defined as pairs of individuals with kinship coefficient < 1/2^(^9^/^2^)^)^10^. After removing individuals who are outliers for genotype heterozygosity and/or missingness, we obtain a set of *N* = 290,641 unrelated British individuals to use in the real data analyses. For all traits, marginal association statistics are computed through OLS in PLINK, using age, sex, and the top 20 genetic principal components (PCs) as covariates in the regression; these 20 PCs were precomputed by UK Biobank from a superset of 488,295 individuals. Additional covariates were used for waist-to-hip ratio (adjusted for BMI) and diastolic/systolic blood pressure (adjusted for cholesterol-lowering medication, blood pressure medication, insulin, hormone replacement therapy, and oral contraceptives). We compute 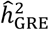 for each trait using chromosome-wide in-sample LD estimated from all *N* individuals.

When using LDSC, S-LDSC, or SumHer to estimate SNP-heritability, it is necessary to define and distinguish between the following sets of SNPs: the set of SNPs containing all possible causal SNPs of interest (used to compute LD scores and LDAK weights), the set of SNPs used as observations in the regression, and the set of SNPs that defines the SNP-heritability estimand of interest. We run two versions of LDSC, S-LDSC (controlling for the most recent baseline-LD model^12,13,30^), and SumHer^14^. First, to enable a more direct comparison between 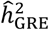 and the estimands of LDSC, S-LDSC, and SumHer, we run an “in-sample LD” version of each method where the *M* typed SNPs (MAF > 0.01) are used to compute LD scores and LDAK weights, perform the regression, and estimate SNP-heritability (i.e. we define the SNP-heritability estimand to be the sum of the per-SNP variances across the *M* typed SNPs). We refer to the in-sample LD versions of these methods as LDSC (in-sample), S-LDSC (baseline-LD/in-sample), and SumHer (in-sample). To run LDSC (in-sample) and S-LDSC (baseline-LD/in-sample), we use the LDSC software (URLs) to compute LD scores and regression weights within 2-Mb windows centered on each SNP, using a random sample of 40K individuals to reduce the memory requirement. To run SumHer (in-sample), we use the LDAK software (URLs) to compute LD tagging from the genotypes of all *N* individuals, using 1-Mb windows centered on each SNP and setting *α* = −0.25 as recommended^9,14^. Unless otherwise specified, all other parameters were set to the default settings of each software.

To enable comparisons between 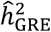 and estimates from LDSC, S-LDSC, and SumHer reported in the literature, we also run each method with its recommended parameter settings and LD estimated from reference panel sequencing data. We refer to these methods as LDSC (1KG), S-LDSC (baseline-LD/1KG), and SumHer (1KG) to indicate that LD is estimated from 489 Europeans in the 1000 Genomes Phase 3 reference panel^31^. We run LDSC (1KG) and S-LDSC (baseline-LD/1KG) with LD scores and regression weights computed within 1-cM windows from 9,997,231 SNPs with minor allele count greater than 5 in the reference panel (URLs), and we define the SNP-heritability estimand to be a function of the array SNPs with MAF > 0.05^11,12^. We run SumHer (1KG) using 8,569,062 SNPs with MAF > 0.01 in the reference panel to compute LDAK weights and LD tagging (1-cM windows) and to define the SNP-heritability estimand; we control for a multiplicative inflation of test statistics as recommended^14^. See refs.^11,12,14,19^ for details about the definitions and interpretations of the estimands of LDSC, S-LDSC, and SumHer.

## References

1. Visscher, P. M., Hill, W. G. & Wray, N. R. Heritability in the genomics era — concepts and misconceptions. Nat. Rev. Genet. 9, 255 (2008).

2. Wray, N. R. et al. Pitfalls of predicting complex traits from SNPs. Nat. Rev. Genet. 14, 507 (2013).

3. Yang, J. et al. Common SNPs explain a large proportion of the heritability for human height. Nat. Genet. 42, 565–569 (2010).

4. Visscher, P. M., Brown, M. A., McCarthy, M. I. & Yang, J. Five Years of GWAS Discovery. Am. J. Hum. Genet. 90, 7–24 (2012).

5. Visscher, P. M. et al. 10 Years of GWAS Discovery: Biology, Function, and Translation. Am. J. Hum. Genet. 101, 5–22 (2017).

6. Speed, D., Hemani, G., Johnson, M. R. & Balding, D. J. Improved Heritability Estimation from Genome-wide SNPs. Am. J. Hum. Genet. 91, 1011–1021 (2012).

7. Yang, J. et al. Genetic variance estimation with imputed variants finds negligible missing heritability for human height and body mass index. Nat. Genet. 47, 1114 (2015).

8. Loh, P.-R. et al. Contrasting genetic architectures of schizophrenia and other complex diseases using fast variance-components analysis. Nat. Genet. 47, 1385 (2015).

9. Speed, D. et al. Reevaluation of SNP heritability in complex human traits. Nat. Genet. 49, 986 (2017).

10. Bycroft, C. et al. The UK Biobank resource with deep phenotyping and genomic data. Nature 562, 203–209 (2018).

11. Bulik-Sullivan, B. K. et al. LD Score regression distinguishes confounding from polygenicity in genome-wide association studies. Nat Genet advance on, 291–295 (2015).

12. Finucane, H. K. et al. Partitioning heritability by functional annotation using genome-wide association summary statistics. Nat Genet 47, 1228–1235 (2015).

13. Gazal, S. et al. Linkage disequilibrium–dependent architecture of human complex traits shows action of negative selection. Nat. Genet. 49, 1421 (2017).

14. Speed, D. & Balding, D. J. SumHer better estimates the SNP heritability of complex traits from summary statistics. Nat. Genet. (2018). doi:10.1038/s41588-018-0279-5

15. Haseman, J. K. & Elston, R. C. The investigation of linkage between a quantitative trait and a marker locus. Behav. Genet. 2, 3–19 (1972).

16. Wu, Y. & Sankararaman, S. A scalable estimator of SNP heritability for biobank-scale data. Bioinformatics 34, i187–i194 (2018).

17. Timpson, N. J., Greenwood, C. M. T., Soranzo, N., Lawson, D. J. & Richards, J. B. Genetic architecture: the shape of the genetic contribution to human traits and disease. Nat. Rev. Genet. 19, 110 (2017).

18. Evans, L. M. et al. Comparison of methods that use whole genome data to estimate the heritability and genetic architecture of complex traits. Nat. Genet. 50, 737–745 (2018).

19. Gazal, S., Marquez-Luna, C., Finucane, H. K. & Price, A. L. Reconciling S-LDSC and LDAK models and functional enrichment estimates. bioRxiv 256412 (2018). doi:10.1101/256412

20. Eyre-Walker, A. Genetic architecture of a complex trait and its implications for fitness and genome-wide association studies. Proc. Natl. Acad. Sci. 107, 1752 LP–1756 (2010).

21. Lohmueller, K. E. The Impact of Population Demography and Selection on the Genetic Architecture of Complex Traits. PLOS Genet. 10, e1004379 (2014).

22. Schoech, A. et al. Quantification of frequency-dependent genetic architectures and action of negative selection in 25 UK Biobank traits. bioRxiv 188086 (2017). doi:10.1101/188086

23. Zeng, J. et al. Signatures of negative selection in the genetic architecture of human complex traits. Nat. Genet. 50, 746–753 (2018).

24. O’Connor, L. J. et al. Polygenicity of complex traits is explained by negative selection. bioRxiv 420497 (2018). doi:10.1101/420497

25. Uricchio, L. H., Kitano, H. C., Gusev, A. & Zaitlen, N. A. An evolutionary compass for detecting signals of polygenic selection and mutational bias. Evol. Lett. 3, 69–79 (2019).

26. Zhang, Y., Qi, G., Park, J.-H. & Chatterjee, N. Estimation of complex effect-size distributions using summary-level statistics from genome-wide association studies across 32 complex traits. Nat. Genet. 50, 1318–1326 (2018).

27. Loh, P.-R., Kichaev, G., Gazal, S., Schoech, A. P. & Price, A. L. Mixed-model association for biobank-scale datasets. Nat. Genet. 50, 906–908 (2018).

28. Gamazon, E. R., Cox, N. J. & Davis, L. K. Structural Architecture of SNP Effects on Complex Traits. Am. J. Hum. Genet. 95, 477–489 (2014).

29. Shi, H., Kichaev, G. & Pasaniuc, B. Contrasting the Genetic Architecture of 30 Complex Traits from Summary Association Data. Am. J. Hum. Genet. 99, 139–153 (2016).

30. Gazal, S. et al. Functional architecture of low-frequency variants highlights strength of negative selection across coding and non-coding annotations. Nat. Genet. 50, 1600–1607 (2018).

31. Consortium, T. 1000 G. P. et al. A global reference for human genetic variation. Nature 526, 68 (2015).

32. Ledoit, O. & Wolf, M. A well-conditioned estimator for large-dimensional covariance matrices. J. Multivar. Anal. 88, 365–411 (2004).

33. Nagai, A. et al. Overview of the BioBank Japan Project: Study design and profile. J. Epidemiol. 27, S2–S8 (2017).

34. Leitsalu, L. et al. Cohort Profile: Estonian Biobank of the Estonian Genome Center, University of Tartu. Int. J. Epidemiol. 44, 1137–1147 (2015).

35. Gaziano, J. M. et al. Million Veteran Program: A mega-biobank to study genetic influences on health and disease. J. Clin. Epidemiol. 70, 214–223 (2016).

36. Pasaniuc, B. & Price, A. L. Dissecting the genetics of complex traits using summary association statistics. Nat. Rev. Genet. 18, 117 (2016).

37. Hormozdiari, F., Kichaev, G., Yang, W.-Y., Pasaniuc, B. & Eskin, E. Identification of causal genes for complex traits. Bioinformatics 31, i206–i213 (2015).

38. Shi, H., Mancuso, N., Spendlove, S. & Pasaniuc, B. Local Genetic Correlation Gives Insights into the Shared Genetic Architecture of Complex Traits. Am. J. Hum. Genet. 101, 737–751 (2017).

39. Yengo, L. et al. Imprint of assortative mating on the human genome. Nat. Hum. Behav. 2, 948–954 (2018).

40. Golan, D., Lander, E. S. & Rosset, S. Measuring missing heritability: inferring the contribution of common variants. Proc. Natl. Acad. Sci. U. S. A. 111, E5272–81 (2014).

41. Weissbrod, O., Flint, J. & Rosset, S. Estimating SNP-Based Heritability and Genetic Correlation in Case-Control Studies Directly and with Summary Statistics. Am. J. Hum. Genet. 103, 89–99 (2018).

42. Elman, R. S., Karpenko, N. & Merkurjev, A. The algebraic and geometric theory of quadratic forms. 56, (American Mathematical Soc., 2008).

43. Lee, S. H., Wray, N. R., Goddard, M. E. & Visscher, P. M. Estimating Missing Heritability for Disease from Genome-wide Association Studies. Am. J. Hum. Genet. 88, 294–305 (2011).

44. Yang, J., Lee, S. H., Goddard, M. E. & Visscher, P. M. GCTA: A Tool for Genome-wide Complex Trait Analysis. Am. J. Hum. Genet. 88, 76–82 (2011).

45. Purcell, S. et al. PLINK: A Tool Set for Whole-Genome Association and Population-Based Linkage Analyses. Am. J. Hum. Genet. 81, 559–575 (2007).

46. Lee, S. H. et al. Estimating the proportion of variation in susceptibility to schizophrenia captured by common SNPs. Nat. Genet. 44, 247 (2012).

47. Lee, S. H. et al. Estimation of SNP Heritability from Dense Genotype Data. Am. J. Hum. Genet. 93, 1151– 1155 (2013).

