## Supplementary Information for "Accurate estimation of SNP-heritability from biobank-scale data irrespective of genetic architecture"

Kangcheng Hou[1,2*], Kathryn S. Burch[3*†], Arunabha Majumdar[1], Huwenbo Shi[3,4], Nicholas Mancuso[1], Yue Wu[5], Sriram Sankararaman[3,5,6,7], and Bogdan Pasaniuc[1,3,6,7,†]

[1]Department of Pathology and Laboratory Medicine, David Geffen School of Medicine, University of California, Los Angeles, Los Angeles, CA, USA
[2]College of Computer Science and Technology, Zhejiang University, Hangzhou, Zhejiang, China
[3]Bioinformatics Interdepartmental Program, University of California, Los Angeles, Los Angeles, CA, USA
[4]Department of Epidemiology, Harvard T.H. Chan School of Public Health, Boston, MA, USA
[5]Department of Computer Science, University of California, Los Angeles, Los Angeles, CA, USA
[6]Department of Human Genetics, David Geffen School of Medicine, University of California, Los Angeles, Los Angeles, CA, USA
[7]Department of Computational Medicine, David Geffen School of Medicine, University of California, Los Angeles, Los Angeles, CA, USA
[*]These authors contributed equally to this work.

April 1, 2019

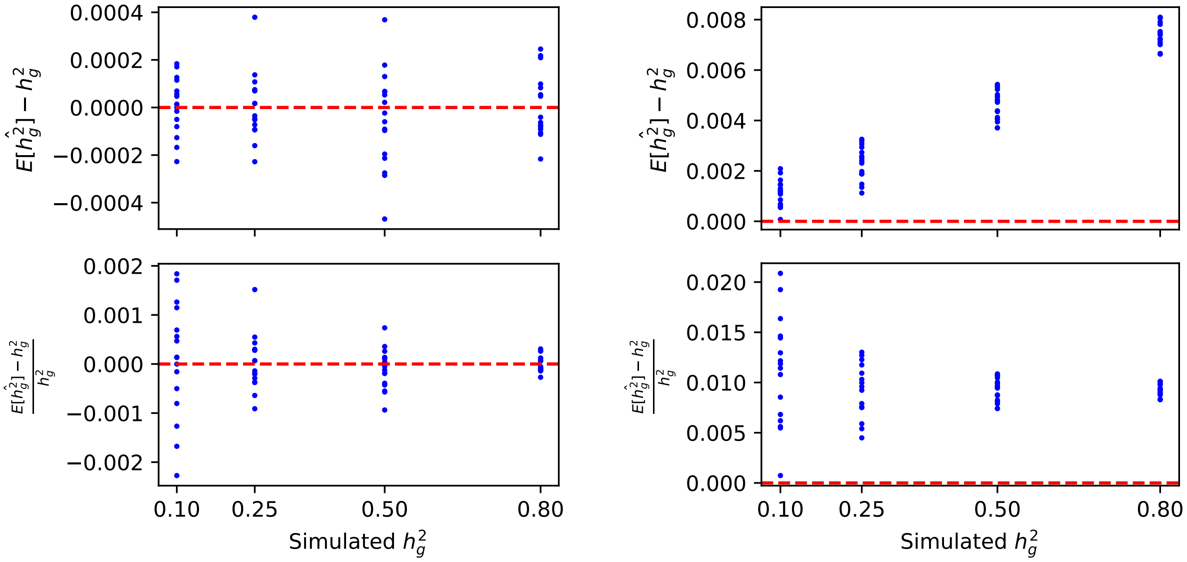


(a) chromosome 22 simulations ($M = 9654$)

(b) genome-wide simulations ($M = 593300$)

Figure S1: Bias and relative bias of $\widehat{h}^2_{\mathrm{GRE}}$ in simulations under 64 MAF- and LDAK-LD-dependent architectures ($N = 337$K). (a) Phenotypes were drawn from $M = 9654$ SNPs on chromosome 22; $h_g^2$ was estimated with a single LD block spanning chromosome 22. (b) Phenotypes were drawn from $M = 593300$ SNPs genome-wide; $h_g^2$ was estimated using 22 chromosome-wide LD blocks. Each point represents the magnitude of the bias of $\widehat{h}^2_{\mathrm{GRE}}$ (top row) or the bias of $\widehat{h}^2_{\mathrm{GRE}}$ relative to the simulated $h_g^2$ (bottom row) estimated from 100 simulations under a single genetic architecture.

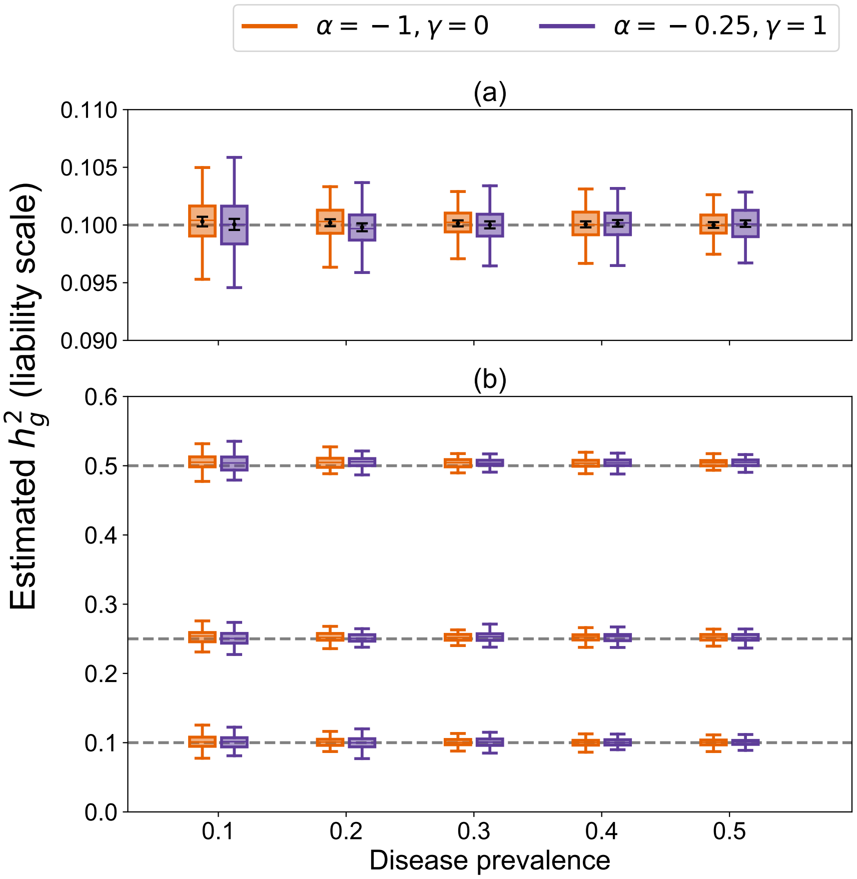


Figure S2: $\widehat{h}^2_{\mathrm{GRE}}$ for case-control GWAS with no ascertainment ($N = 337$K). Each boxplot represents estimates from 100 independent simulations at the specified disease prevalence. In all simulations, $p_{\mathrm{causal}} = 1$ and causal variants were drawn uniformly. (a) Each individual's liability was drawn from $M = 9654$ SNPs on chromosome 22 and converted to a binary case-control status; $h_g^2$ was estimated with a single block. Black points and error bars represent the mean and $\pm 2$ s.e.m. (b) Each individual's liability was drawn from $M = 593300$ SNPs genome-wide and converted to a binary case-control status; $h_g^2$ was estimated with 22 chromosome-wide LD blocks.

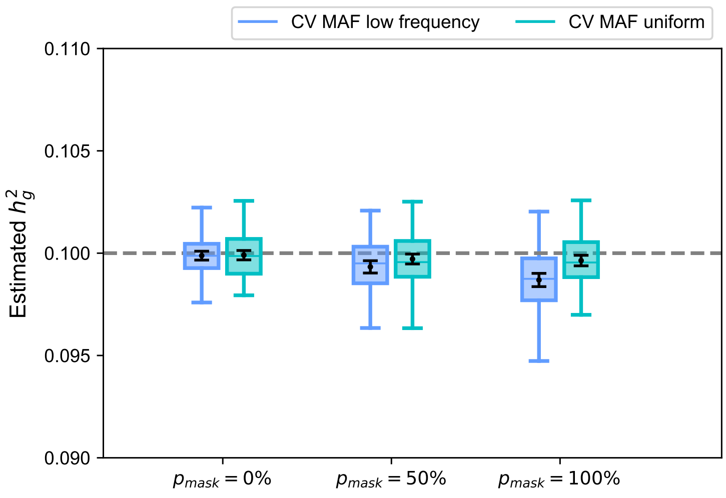


Figure S3: $\widehat{h}^2_{\mathrm{GRE}}$ in simulations on chromosome 22 where a percentage of causal SNPs are masked from the observed summary statistics ($p_{\mathrm{mask}} = 0\%$, $50\%$, or $100\%$). "CV MAF low frequency" refers to CV MAF $= [0.01, 0.05]$. "CV MAF uniform" means causal variants were drawn uniformly from the chromosome 22 typed SNPs.

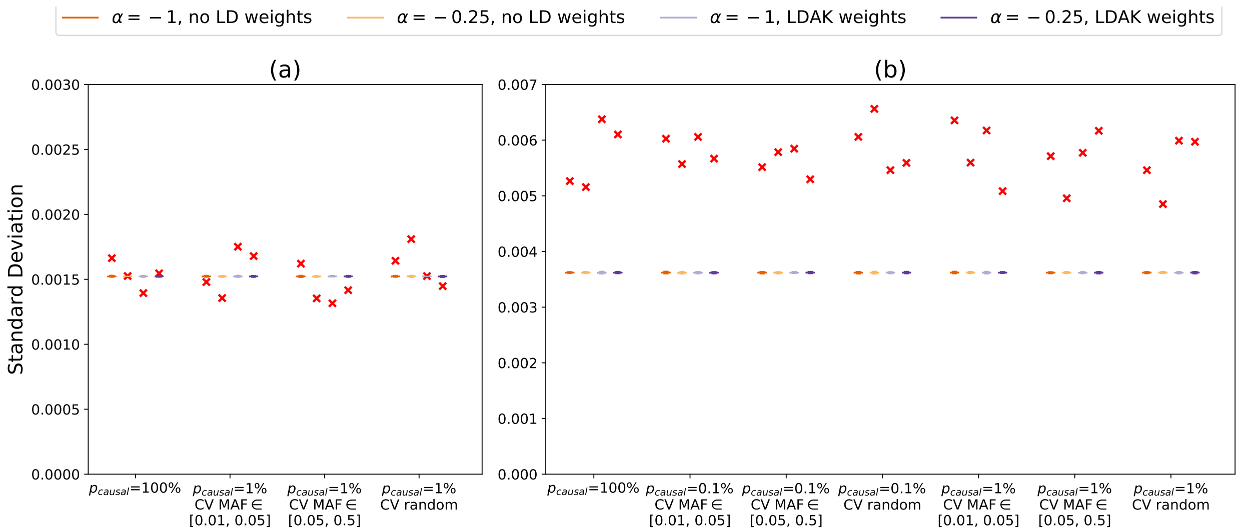


Figure S4: Comparison of the analytical standard error of $\widehat{h}^2_{\mathrm{GRE}}$ with the standard deviation of $\widehat{h}^2_{\mathrm{GRE}}$ computed from 100 simulations ($h_g^2 = 0.25$). (a) Phenotypes were simulated from SNPs on chromosome 22 ($N = 337205$, $M = 9564$ array SNPs) under one of 16 LDAK-LD- and/or MAF-dependent architectures and $\widehat{h}^2_{\mathrm{GRE}}$ was computed with a single chromosome-wide LD block. (b) Phenotypes were simulated from all genome-wide SNPs ($N = 337205$, $M = 593300$ array SNPs) under one of 28 LDAK-LD- and/or MAF-dependent architectures and $\widehat{h}^2_{\mathrm{GRE}}$ was computed with 22 chromosome-wide LD blocks. The colored bars represent the distribution of standard error estimates from 100 simulations. The red crosses mark the empirical standard deviation of the 100 estimates of $h_g^2$.

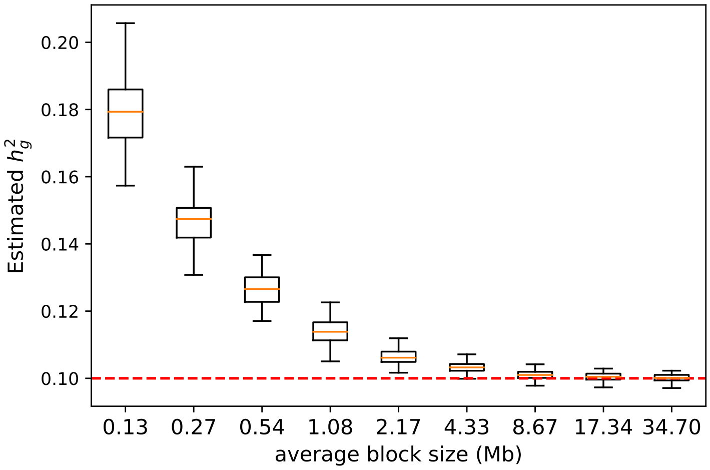


Figure S5: Distribution of $\widehat{h}^2_{\text{GRE}}$ in simulations on chromosome 22 ($N = 337205$, $M = 9564$ array SNPs) as a function of the average size (Mb) of the LD blocks that were used to compute $\widehat{h}^2_{\text{GRE}}$. The largest block size (34.70 Mb) corresponds to using a single chromosome-wide LD block. All simulations were performed $h_g^2 = 0.1$, $p_{\text{causal}} = 0.01$, $\alpha = -1$, and $\gamma = 0$ (no LD weights). Each boxplot represents 100 estimates.

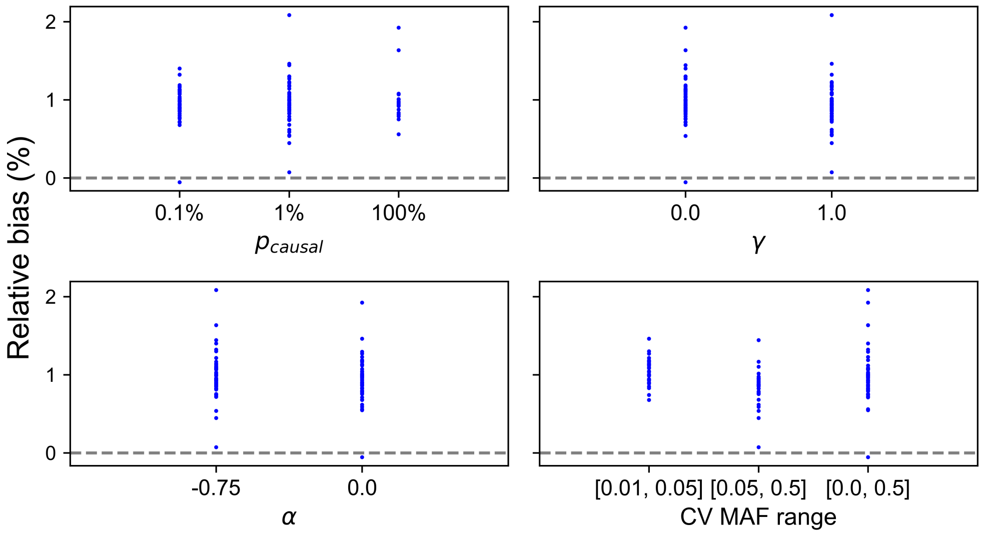


Figure S6: Relative bias of $\widehat{h}^2_{\text{GRE}}$ in genome-wide simulations ($N = 337\text{K}$, $M = 593\text{K}$) with respect to different values of $p_{\text{causal}}$, $\alpha$, $\gamma$, and CV MAF. Each point is the estimated relative bias of GRE (as a percentage of the simulated $h_g^2$) for a single architecture. Each plot contains the results from the same 64 architectures shown in Figure 1b and Supplementary Table S1b.

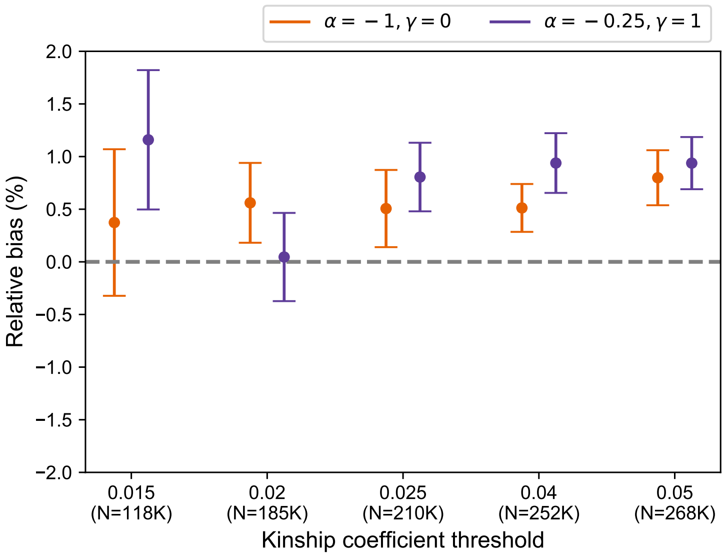


Figure S7: Relative bias of $\widehat{h}^2_{\text{GRE}}$ in genome-wide simulations ($M = 593\text{K}$) in which individuals were filtered at different kinship coefficient thresholds. Kinship matrix is defined as $\mathbf{X}\mathbf{X}^T/M$. Each point marks the relative bias (as a percentage of $h_g^2$) estimated from 100 independent simulations; bars represent $\pm 2$ s.e.m. In all simulations, $h_g^2 = 0.25$, $p_{\text{causal}} = 1$, causal variants are drawn uniformly, and $\widehat{h}^2_{\text{GRE}}$ is computed with 22 chromosome-wide blocks.

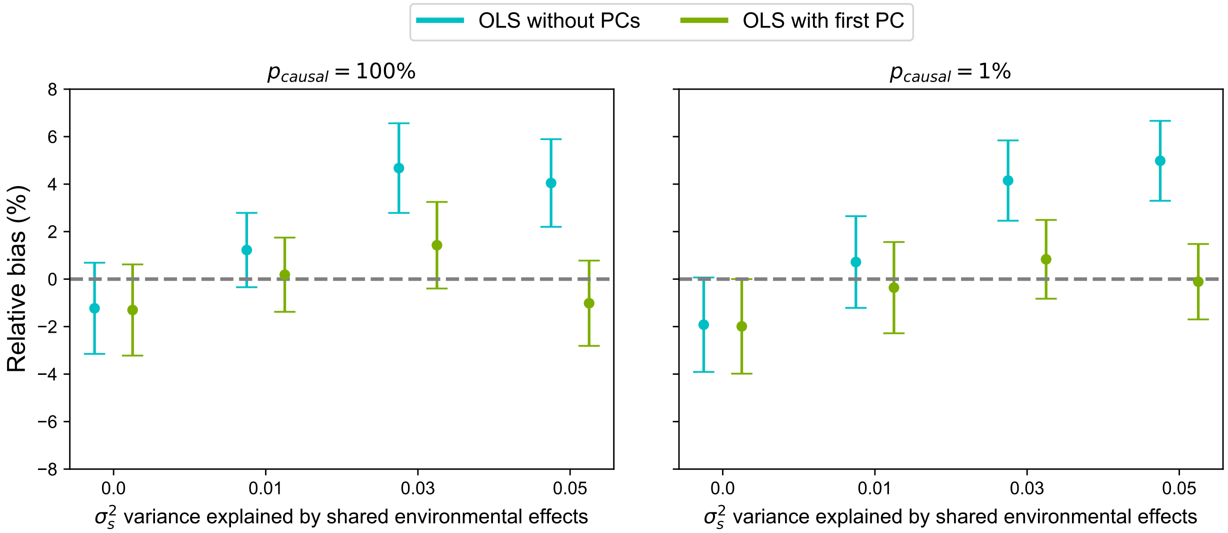


Figure S8: Relative bias of $\widehat{h}^2_{\text{GRE}}$ in genome-wide simulations ($N = 8430$, $M = 14821$) with population stratification (see Methods). $\sigma_s^2$ is the proportion of total phenotypic variance explained by the covariate (i.e. the first genetic PC). Other simulation parameters are fixed ($h_g^2 = 0.25$, $\alpha = -1$, $\gamma = 0$) and causal variants are drawn uniformly.

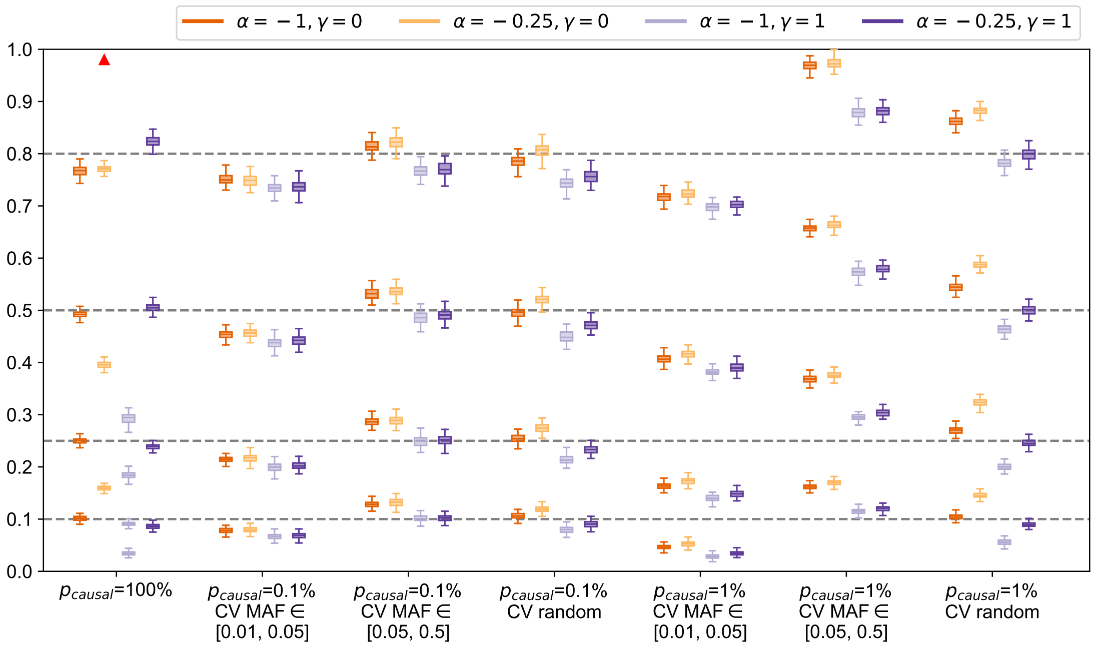


Figure S9: Distribution of $h_g^2$ estimates from LDSC (no annotations) in simulations across 112 LDAK-LD- and/or MAF-dependent architectures ($N$ = 337205 individuals, $M$ = 593300 array SNPs). Each boxplot represents the distribution of 100 estimates under a single architecture. Boxplot whiskers extend to the minimum and maximum estimates located within 1.5 × IQR from the first and third quartiles, respectively.

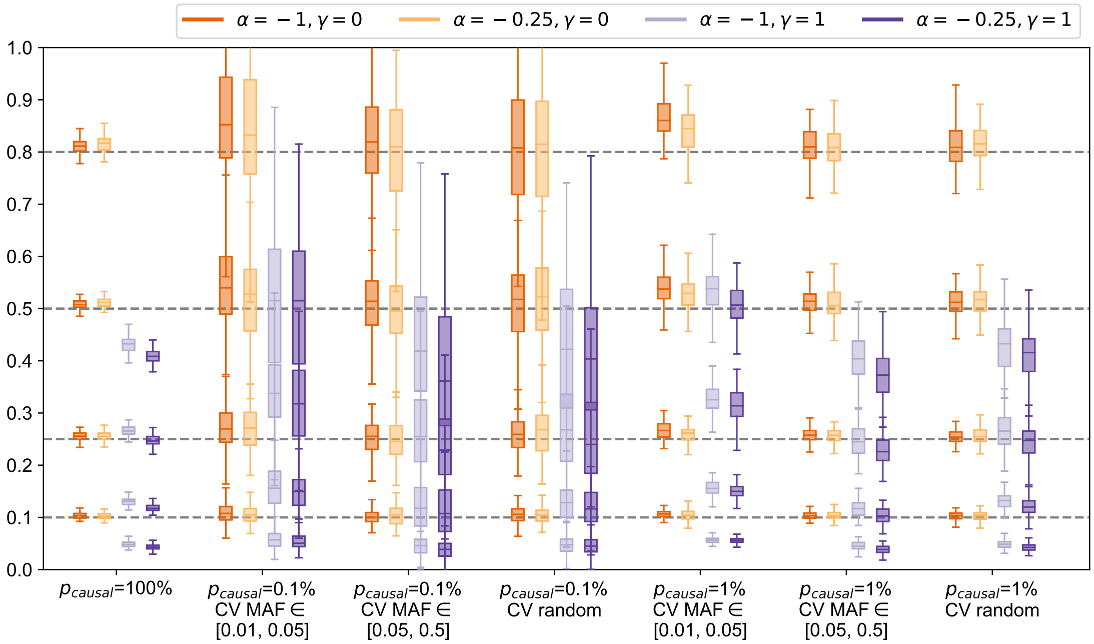


Figure S10: Distribution of $h_g^2$ estimates from S-LDSC (10 MAF bins) in simulations across 112 LDAK-LD- and/or MAF-dependent architectures ($N$ = 337205 individuals, $M$ = 593300 array SNPs). Each boxplot represents the distribution of 100 estimates under a single architecture. Boxplot whiskers extend to the minimum and maximum estimates located within 1.5 × IQR from the first and third quartiles, respectively.

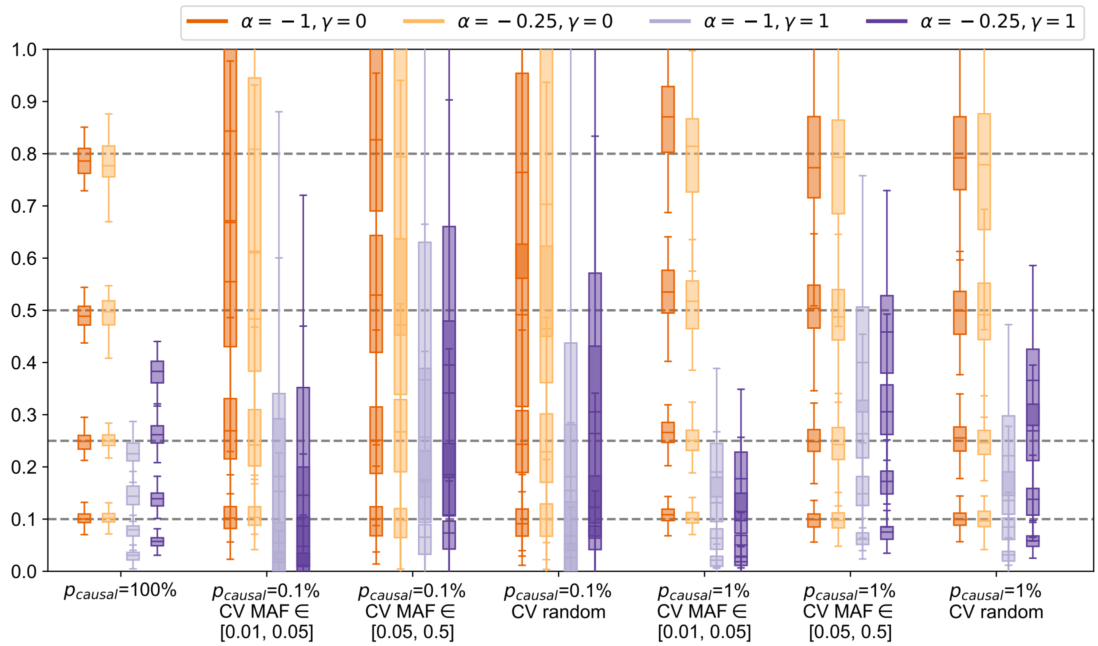


Figure S11: Distribution of $h_g^2$ estimates from S-LDSC (10 MAF bins + LLD) in simulations across 112 LDAK-LD- and/or MAF-dependent architectures ($N = 337205$ individuals, $M = 593300$ array SNPs). Each boxplot shows the distribution of 100 estimates under a single architecture. Boxplot whiskers extend to the minimum and maximum estimates located within 1.5 × IQR from the first and third quartiles, respectively.

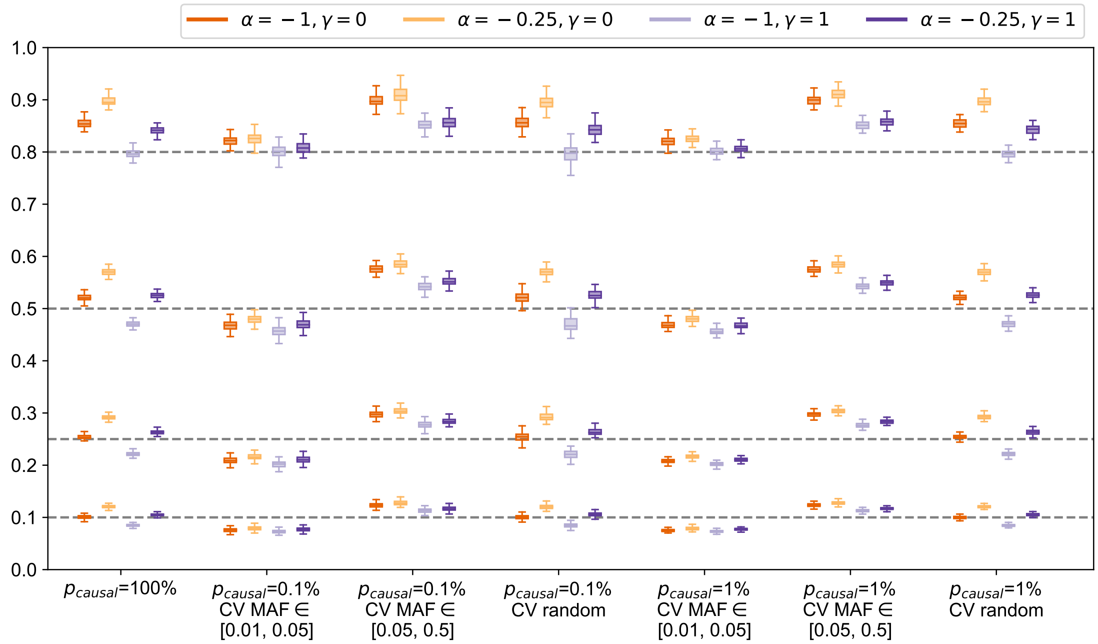


Figure S12: Distribution of $h_g^2$ estimates from SumHer in simulations across 112 LDAK-LD- and/or MAF-dependent architectures ($N = 337205$ individuals, $M = 593300$ array SNPs). Each boxplot represents the distribution of 100 estimates under a single architecture. Boxplot whiskers extend to the minimum and maximum estimates located within 1.5 × IQR from the first and third quartiles, respectively.

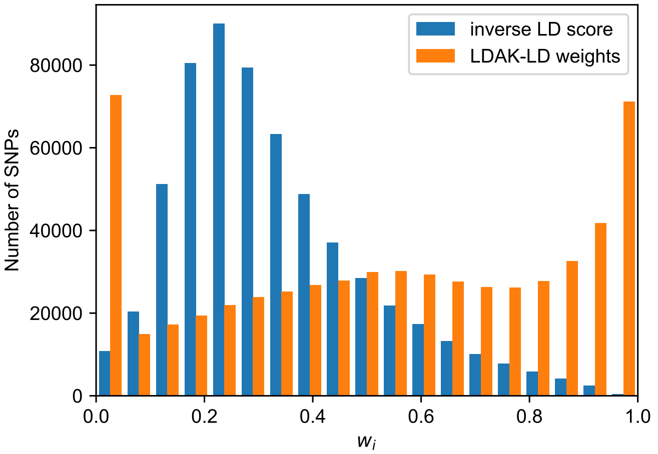


Figure S13: Histograms of LDAK weights and inverse LD score weights used in genome-wide simulations ($M = 593$K SNPs).

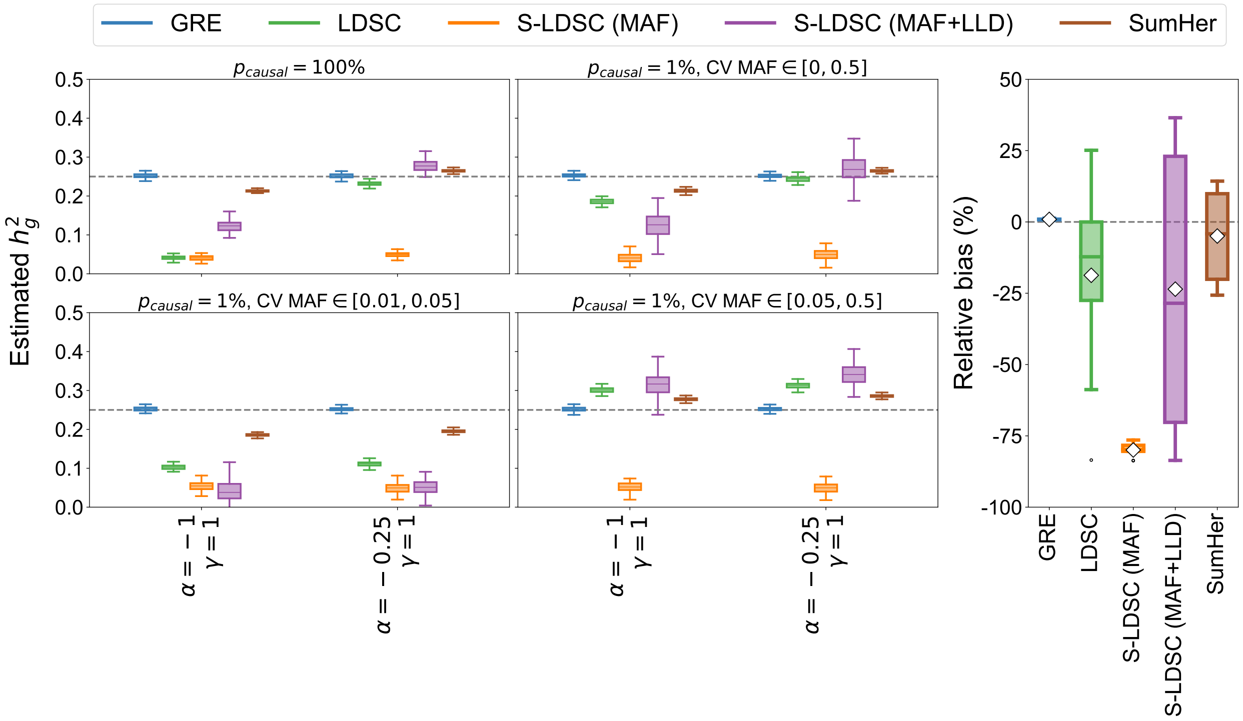


Figure S14: Comparison of methods across 14 MAF- and LD-score-dependent architectures ($N = 337205$ individuals, $M = 593300$ array SNPs, $h_g^2 = 0.25$). LD-score-dependent architectures are simulated by coupling the variance of each SNP to the inverse of its LD score (Methods). **Left:** Each boxplot represents 100 estimates under a single architecture; results are shown for $p_{\text{causal}} = 100\%$ and $1\%$. **Right:** Each boxplot represents the distribution of the relative bias across all 14 LD-score-dependent architectures. White diamonds mark the average of each distribution. All boxplot whiskers mark the minimum and maximum estimates located within $1.5 \times$ IQR from the first and third quartiles, respectively.

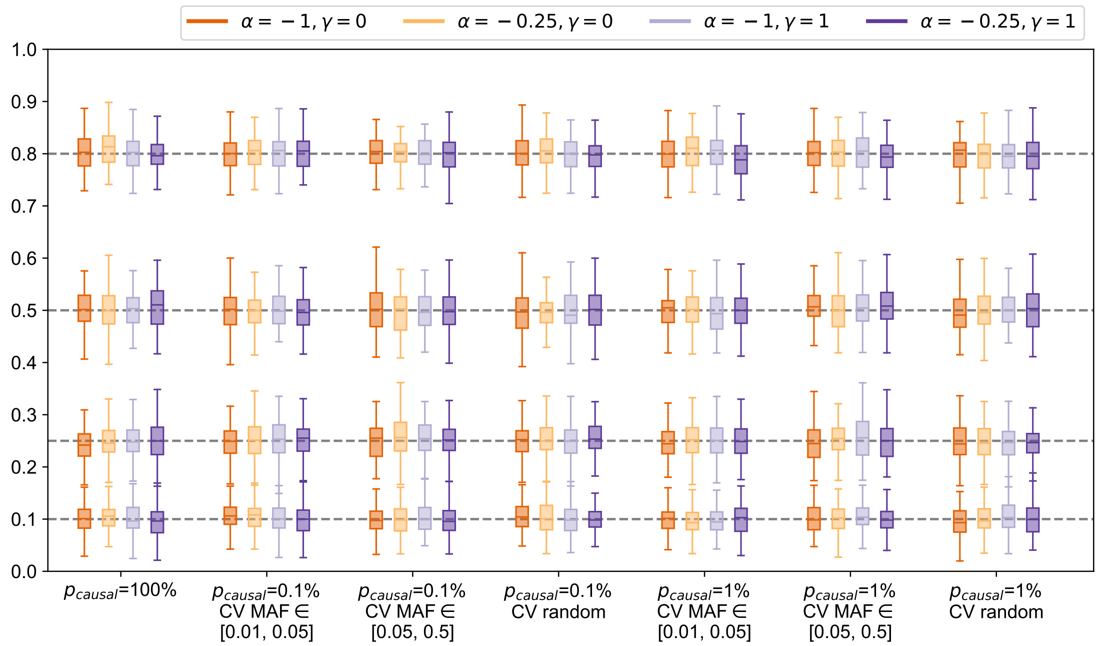


Figure S15: Distribution of $h_g^2$ estimates from GRE in simulations across 112 LDAK-LD- and/or MAF-dependent architectures ($N$ = 8430 individuals, $M$ = 14821 array SNPs). Each boxplot represents the distribution of 100 estimates under a single architecture. Boxplot whiskers extend to the minimum and maximum estimates located within 1.5 × IQR from the first and third quartiles, respectively.

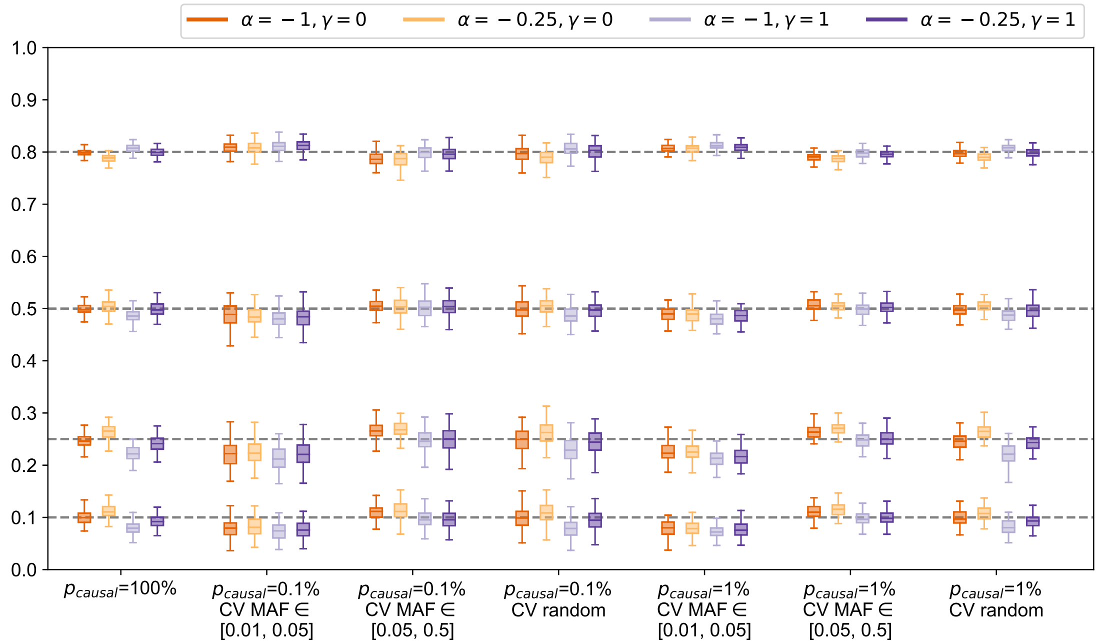


Figure S16: Distribution of $h_g^2$ estimates from single-component GREML in simulations across 112 LDAK-LD- and/or MAF-dependent architectures ($N$ = 8430 individuals, $M$ = 14821 array SNPs). Each boxplot shows the distribution of 100 estimates under a single architecture. Boxplot whiskers extend to the minimum and maximum estimates located within 1.5 × IQR from the first and third quartiles, respectively.

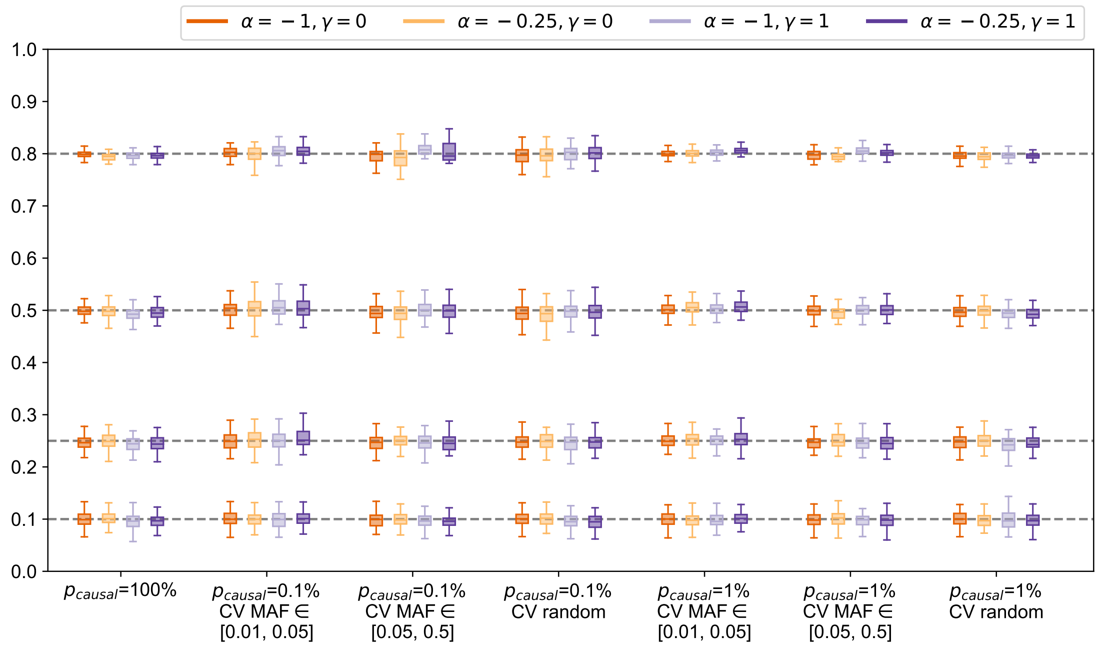


Figure S17: Distribution of $h_g^2$ estimates from GREML-LDMS-I in simulations across 112 LDAK-LD- and/or MAF-dependent architectures ($N$ = 8430 individuals, $M$ = 14281 array SNPs). Each boxplot shows the distribution of 100 estimates under a single architecture. Boxplot whiskers extend to the minimum and maximum estimates located within 1.5 × IQR from the first and third quartiles, respectively.

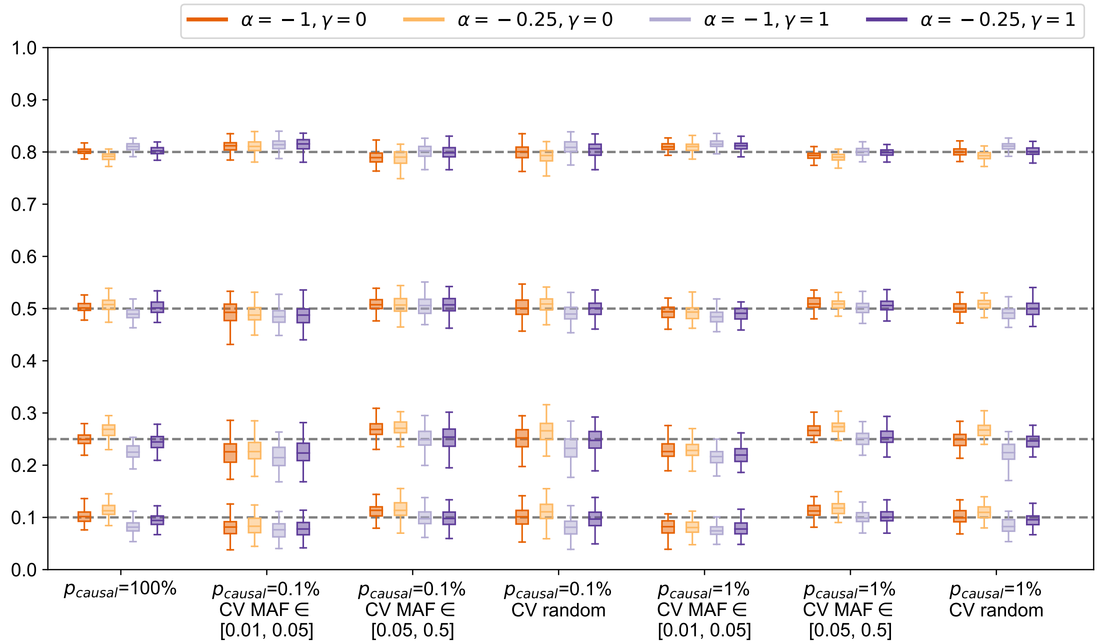


Figure S18: Distribution of $h_g^2$ estimates from BOLT-REML in simulations across 112 LDAK-LD- and/or MAF-dependent architectures ($N$ = 8430 individuals, $M$ = 14281 array SNPs). Each boxplot represents the distribution of 100 estimates under a single architecture. Boxplot whiskers extend to the minimum and maximum estimates located within 1.5 × IQR from the first and third quartiles, respectively.

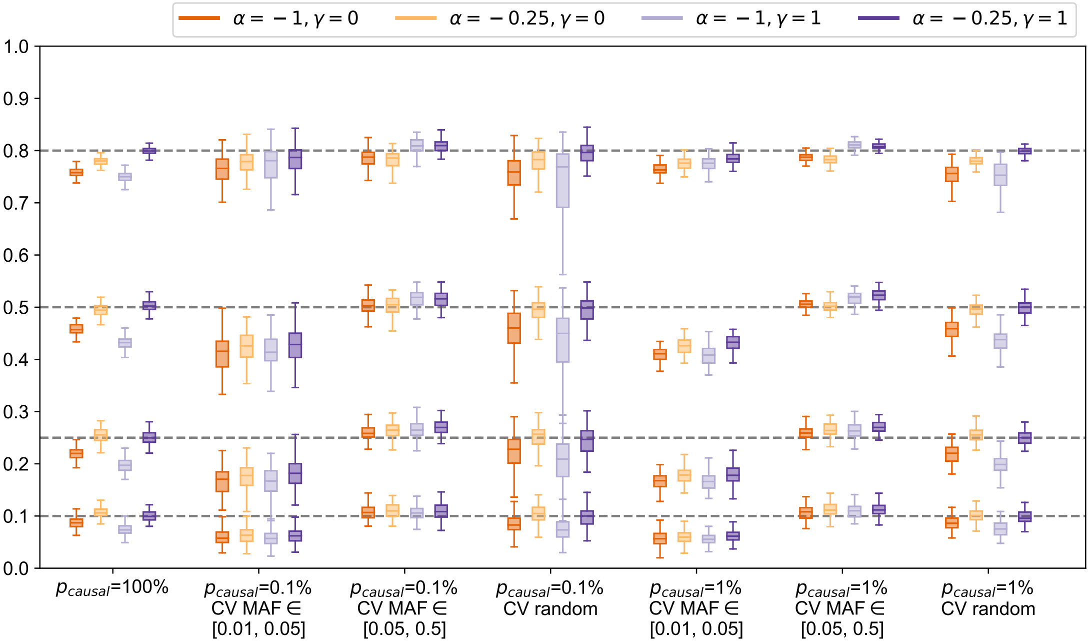


Figure S19: Distribution of $h_g^2$ estimates from LDAK in simulations across 112 LDAK-LD- and/or MAF-dependent architectures ($N = 8430$ individuals, $M = 14281$ array SNPs). Each boxplot represents the distribution of 100 estimates under a single architecture. Boxplot whiskers extend to the minimum and maximum estimates located within 1.5 × IQR from the first and third quartiles, respectively.

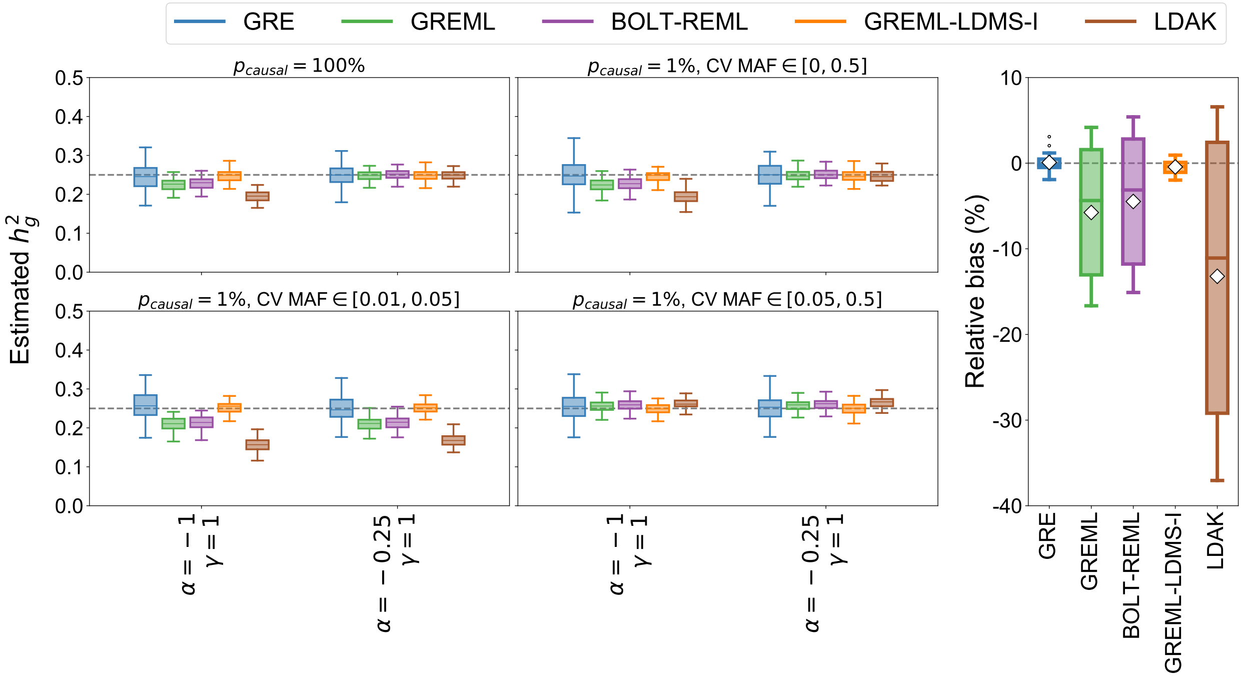


Figure S20: Comparison of methods across 14 MAF- and LD-score-dependent architectures ($N = 8430$ individuals, $M = 14281$ array SNPs). LD-score-dependent architectures are simulated by coupling the variance of each SNP to the inverse of its LD score (see Methods). **Left:** Each boxplot represents 100 estimates under a single architecture; results are shown for $p_{\text{causal}} = 100\%$ and 1%. **Right:** Each boxplot represents the distribution of the relative bias across all 14 LD-score-dependent architectures. White diamonds mark the average of each distribution. All boxplot whiskers mark the minimum and maximum estimates located within 1.5 × IQR from the first and third quartiles, respectively.

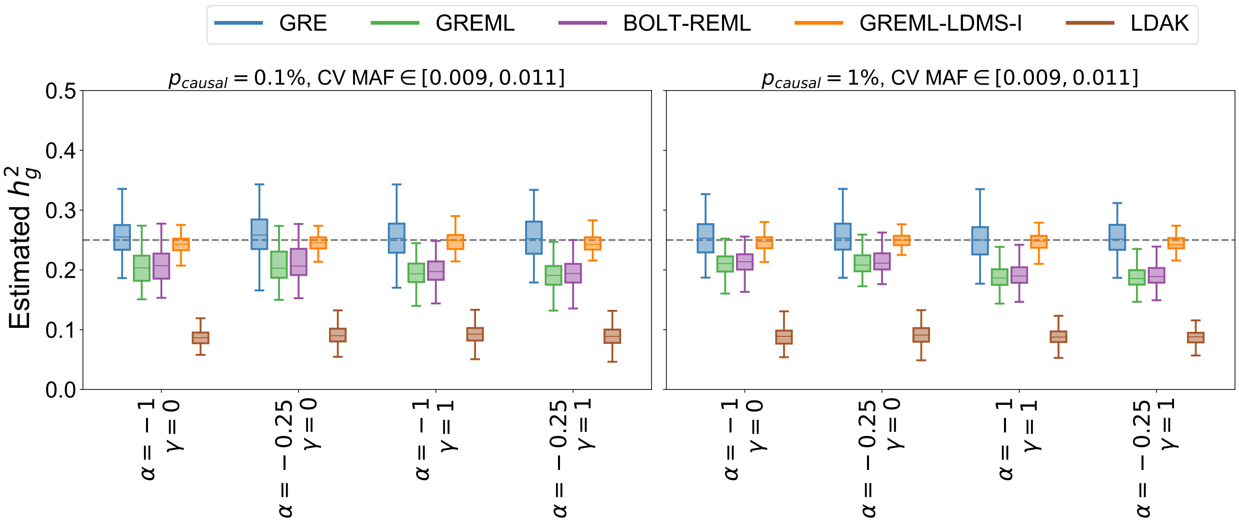


Figure S21: Comparison of GRE, GREML, BOLT-REML, GREML-LDMS-I, and LDAK in small-scale simulations ($N = 8430$ individuals, $M = 14821$ array SNPs) under MAF- and/or LDAK-LD-dependent architectures where all causal variants were drawn from the MAF range $[0.009, 0.011]$. Each boxplot contains estimates of $h_g^2$ from 100 simulations. The GRE estimator was computed with 22 chromosome-wide LD blocks. For GREML-LDMS-I, 8 GRMs were used (2 MAF bins $\times$ 4 LD quartiles). Boxplot whiskers mark the minimum and maximum estimates located within 1.5$\times$ IQR units from the first and third quartiles, respectively.

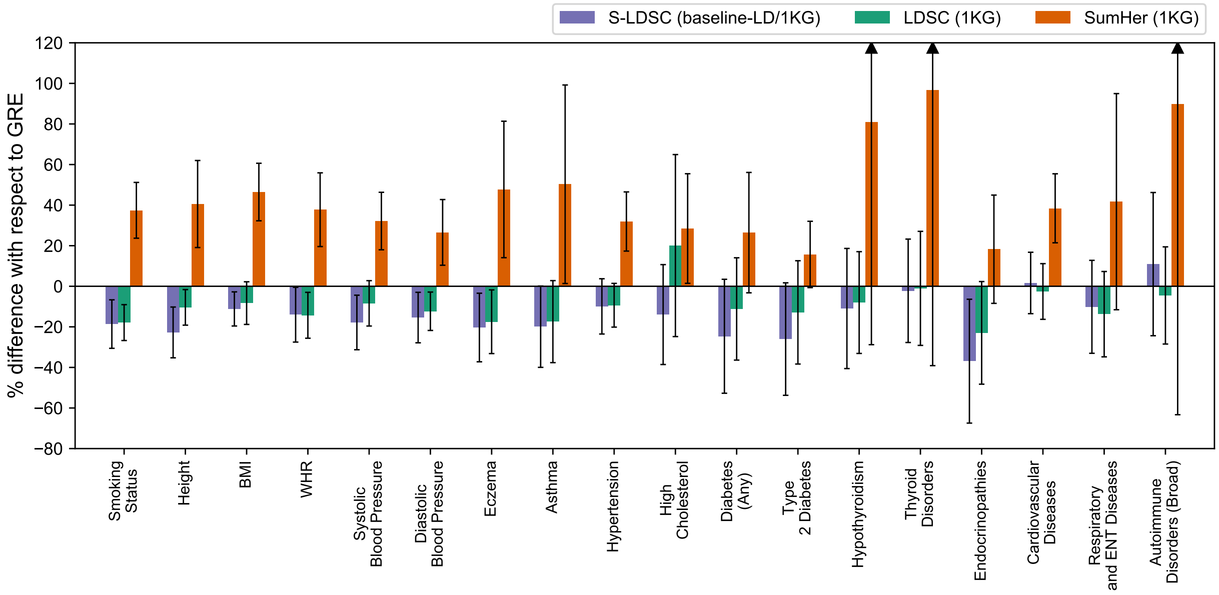


Figure S22: Percent difference of SNP-heritability estimates from LDSC (1KG), S-LDSC (baseline-LD/1KG), and SumHer (1KG) with respect to $\widehat{h}^2_{\mathrm{GRE}}$ for 18 complex traits and diseases in the UK Biobank for which $\widehat{h}^2_{\mathrm{GRE}} > 0.05$ ($N = 290$K unrelated British individuals and $M = 460$K typed SNPs; see Methods). Each bar represents the difference between the estimated SNP-heritability and $\widehat{h}^2_{\mathrm{GRE}}$ as a percentage of $\widehat{h}^2_{\mathrm{GRE}}$. Black bars mark $\pm 2$ standard errors.

### Supplementary Tables (see Excel file)

Table S1: Summary of $\widehat{h}^2_{\text{GRE}}$ in chromosome 22 simulations of a quantitative trait ($N = 337205$ individuals, $M = 9654$ array SNPs, 1 chromosome-wide LD block).

Table S2: Summary of $\widehat{h}^2_{\text{GRE}}$ in chromosome 22 simulations of a case-control GWAS with no ascertainment ($N = 337205$, $M = 9654$, 1 chromosome-wide LD block, $h^2_g = 0.1$). Genetic architecture is either "GREML-SC" ($\alpha = -1$ and $\gamma = 0$) or "LDAK" ($\alpha = -0.25$ and $\gamma = 1$). $p = \{0.1, 0.2, 0.3, 0.4, 0.5\}$ is the population disease prevalence.

Table S3: Summary of $\widehat{h}^2_{\text{GRE}}$ in chromosome 22 simulations of an ascertained case-control study ($N_{\text{case}} = N_{\text{control}} = 33720$, $M = 9654$, $h^2_g = 0.1$, disease prevalence = 0.1). Genetic architecture is either "GREML-SC" ($\alpha = -1$ and $\gamma = 0$) or "LDAK" ($\alpha = -0.25$ and $\gamma = 1$).

Table S4: Comparison of analytical standard error of $\widehat{h}^2_{\text{GRE}}$ and empirical standard deviation of 100 $\widehat{h}^2_{\text{GRE}}$ estimates in simulations on chromosome 22.

Table S5: Summary of $\widehat{h}^2_{\text{GRE}}$ in chromosome 22 simulations ($N = 337205$ individuals, $M = 9654$ array SNPs) where the number of blocks $K$ was varied. Phenotypes were simulated for $h^2_g = 0.1$, $p_{causal} = 0.01$, $\alpha = -1$, and no LD weights ($\gamma = 0$).

Table S6: Summary of $\widehat{h}^2_{\text{GRE}}$ in genome-wide simulations of a quantitative trait ($N = 337205$ individuals, $M = 593300$ array SNPs, 22 chromosome-wide LD blocks).

Table S7: Summary of $\widehat{h}^2_{\text{GRE}}$ in genome-wide simulations of an unascertained case-control study ($N = 337205$ individuals, $M = 593300$ array SNPs, 22 chromosome-wide LD blocks). Genetic architecture is either "GREML-SC" ($\alpha = -1$ and $\gamma = 0$) or "LDAK" ($\alpha = -0.25$ and $\gamma = 1$). $p = \{0.1, 0.2, 0.3, 0.4, 0.5\}$ is the population disease prevalence.

Table S8: Summary of $\widehat{h}^2_{\text{GRE}}$ in simulations using genotypes of $N = 7685$ South Asian individuals and $M = 1642$ SNPs (a subset of SNPs on chr21 and chr22). $K = 1$ means a single LD matrix was used; $K = 2$ means that two "chromosome-wide" LD blocks were used.

Table S9: Comparison of analytical standard error of $\widehat{h}^2_{\text{GRE}}$ and empirical standard deviation of 100 $\widehat{h}^2_{\text{GRE}}$ estimates in genome-wide simulations.

Table S10: Relative bias of $\widehat{h}^2_{\text{GRE}}$ in genome-wide simulations ($M = 593$K) at different kinship coefficient thresholds. For each kinship threshold, we select a subset of the 337K white British individuals for whom all entries in $\mathbf{X}\mathbf{X}^T/M$ are $\leq$ kinship coefficient threshold.

Table S11: Relative bias of $\widehat{h}^2_{\text{GRE}}$ in genome-wide simulations ($M = 593$K, $N = 337$K) with population stratification. The column var_s refers to $\sigma_s^2$, the amount of phenotypic variance explained by the first genetic PC (see Methods). In all simulations, $h_g^2 = 0.25$, $\alpha = -1$, $\gamma = 0$, and causal variants are drawn uniformly. OLS summary statistics were computed with either no covariates or with the first genetic PC as a covariate.

Table S12: Summary of LDSC results in genome-wide simulations.

Table S13: Summary of SumHer results in genome-wide simulations.

Table S14: Boundaries of 10 MAF bins for S-LDSC (MAF).

Table S15: Summary of S-LDSC (MAF) results in genome-wide simulations.

Table S16: Summary of S-LDSC (MAF+LLD) results in genome-wide simulations.

Table S17: GRE, LDSC, SumHer, S-LDSC (MAF), S-LDSC (MAF+LLD) in genome-wide simulations with LD-score-dependent architectures.

Table S18: Summary of GREML in small-scale simulations.

Table S19: Summary of BOLT-REML in small-scale simulations.

Table S20: Summary of LDAK in small-scale simulations.

Table S21: Summary of GRE in small-scale simulations.

Table S22: Summary of GREML-LDMS-I in small-scale simulations.

Table S23: GRE, GREML, BOLT-REML, GREML-LDMS-I, LDAK in small-scale simulations with LD-score-dependent architectures.

Table S24: $\widehat{h}^2_{\text{GRE}}$ (and standard error) computed at a range of kinship coefficient thresholds for 22 complex traits in the UK Biobank. For each kinship threshold (tol), we excluded individuals with kinship $\geq$ tol with any other individual in the data.

Table S25: $\widehat{h}^2_{\text{GRE}}$ (and standard error) computed from SNPs at three MAF thresholds. Percent relative change is $[\widehat{h}^2_{\text{GRE}}(\text{MAF} > f) - \widehat{h}^2_{\text{GRE}}(\text{MAF} > 0.01)] \times 100 \ / \ \widehat{h}^2_{\text{GRE}}(\text{MAF} > 0.01)$ where $\widehat{h}^2_{\text{GRE}}(\text{MAF} > f)$ is the GRE estimate computed from SNPs with MAF $> f$. (Note: the differences between $\widehat{h}^2_{\text{GRE}}(\text{MAF} > 0.01)$ in this table and the results in Table 2 are due to differences in filtering: here, MAF was computed from the array SNPs whereas in Table 2 (and elsewhere), MAF was computed from the imputed UK Biobank genotype data.)

Table S26: Estimates of $h_g^2$ from the GRE approach, LDSC (in-sample), S-LDSC (baseline-LD/in-sample), SumHer (in-sample), LDSC (1KG), S-LDSC (baseline-LD/1KG), and SumHer (1KG) for 22 complex traits and diseases in the UK Biobank (N = 290K unrelated British individuals, M = 460K typed SNPs). Previously reported BOLT-REML estimates (Loh et al. 2018a) are also included.
